# Mutation rates and selection on synonymous mutations in SARS-CoV-2

**DOI:** 10.1101/2021.01.14.426705

**Authors:** Nicola De Maio, Conor R. Walker, Yatish Turakhia, Robert Lanfear, Russell Corbett-Detig, Nick Goldman

## Abstract

The COVID-19 pandemic has seen an unprecedented response from the sequencing community. Leveraging the sequence data from more than 140,000 SARS-CoV-2 genomes, we study mutation rates and selective pressures affecting the virus. Understanding the processes and effects of mutation and selection has profound implications for the study of viral evolution, for vaccine design, and for the tracking of viral spread. We highlight and address some common genome sequence analysis pitfalls that can lead to inaccurate inference of mutation rates and selection, such as ignoring skews in the genetic code, not accounting for recurrent mutations, and assuming evolutionary equilibrium. We find that two particular mutation rates, G→U and C→U, are similarly elevated and considerably higher than all other mutation rates, causing the majority of mutations in the SARS-CoV-2 genome, and are possibly the result of APOBEC and ROS activity. These mutations also tend to occur many times at the same genome positions along the global SARS-CoV-2 phylogeny (i.e., they are very homoplasic). We observe an effect of genomic context on mutation rates, but the effect of the context is overall limited. While previous studies have suggested selection acting to decrease U content at synonymous sites, we bring forward evidence suggesting the opposite.

## 1 Introduction

The abundant and rapid availability of viral genomic data has had a profound effect on the response to the COVID-19 pandemic, from tracking and tracing of transmission [1, 2, 3], to vaccine and drug development [4]. Genomic SARS-CoV-2 data also allows us to investigate the evolutionary dynamics of the virus such as its mutational and selective pressures [5]. Understanding the contribution of mutation and selection in shaping SARS-CoV-2 genome evolution is important, for example, for drug and vaccine development [6], for predicting variants of clinical and epidemiological importance [7, 8, 9, 10, 11], for understanding the biological mechanisms underlying the virus’ genome evolution (such as recombination [12] and mutagenic immune system responses [13, 14, 15, 16]), to improve the accuracy of phylogenetic approaches for epidemiological applications [17, 18, 19] and for inferring its origin [20].

When deciphering genome evolution, one has to disentangle the effect of mutation and selection affecting the emergence and spread of genetic variants. It has been observed by several studies that SARS-CoV-2 presents a very skewed mutational spectrum, with most observed genetic variation resulting from C→U mutations [21, 5, 22, 23]. It is important to account for these mutational skews when inferring selection, since recurrent mutations can generate a phylogenetic signal that can be confused with positive selection when using phylogenetic methods (e.g. [24, 25]). Such strong mutational skews, if unaccounted for, can also cause errors in phylogenetic tree inference [23, 22].

Here, we identify common pitfalls when analysing SARS-CoV-2 genomic data, and we present alternative, more robust approaches for identifying the contribution of mutation and selection in SARS-CoV-2 evolution. We confirm that the C→U mutation rate is very high in SARS-CoV-2, in particular within the context UCG→UUG, putatively as the result of APOBEC (“Apolipoprotein B mRNA Editing Catalytic Polypeptide-like”) proteins activity. However, the majority of C→U mutations occur outside of this context. Secondly, and in contrast to most other studies, we find that the G→U mutation rate is nearly as high (about 97%) as C→U. The reason why this has not been noted in most other studies is because they did not account for the biases in the genetic code that causes most G→U mutations to be selectively deleterious, particularly compared to C→U mutations. This has a large effect on inferred mutation rates also because most of the SARS-CoV-2 genome is composed of coding sequence. This causes G→U mutations to be underrepresented among the observed genetic variation, despite its high mutation rate. Finally, we investigate selection acting on synonymous variants. We find evidence contrary to previous claims of selection against U content [5].

## 2 New Approaches

Estimating accurate mutation rates is essential for understanding the evolutionary and immunological pressures acting on the virus, as well as to infer accurate phylogenies and for detecting selection. One of the main aims of our work is the estimation of mutation rates in SARS-CoV-2, in particular while trying to account for the effects of selection, which can affect the spread of certain types of mutation, and therefore decrease or increase our chances of detecting them.

Previous studies investigating mutation rates in SARS-CoV-2 have used the approach of counting the number of genome positions at which alternative alleles are observed (see e.g. [5, 21]). This approach works well when the number of mutations occurred is small relative to the number of genome positions considered. However, when the same mutation events occur multiple times at the same position on different branches of the phylogenetic tree (as the case for SARS-CoV-2 [22, 23, 9]), this approach can underestimate the most elevated mutation rates, since multiple mutation events can end up conflated and counted as a single variant allele.

Our approach consists instead of, first, inferring a maximum likelihood phylogenetic tree for considered SARS-CoV-2 genomes. Then, we infer a mutational history on this tree for each position of the genome using parsimony. This gives us, for each given mutation type (for example, synonymous mutations from nucleotide A to C) an estimated number of mutation events. We then normalize this number of mutation events, dividing it by the number of mutation “possibilities”. These possibilities represent the number of sites in the reference genome in which a mutation of a certain type is possible. At each position, multiple mutation events of the same type are possible along the phylogeny, so the inferred number of mutation events of a certain type can be larger than the number of mutation possibilities for the same type. These normalized counts represents our estimates of relative neutral mutation rates.

In order to obtain multiple independent estimates, and in order to assess and possibly correct for the contribution of possible issues in our sequences, we separate mutation counts in 3 bins according to the number of descendants of each mutation: (i) those with only one descendant tip (“singletons”); (ii) those with ≥ 2 and ≤ 4 descendants (“low frequency”); and (iii) those with > 4 descendants (“high frequency”). The main rationale for this is that sequence errors and RNA degradation, for example, are expected to mostly cause apparent mutation events with only 1 or very few descendants.

For completeness, we also show results from the alternative approach to estimating mutation rates used in previous studies, based on counting the number of alignment columns at which alternative alleles are observed. We note again, however, that due to possibly multiple mutation events of the same type occurring at the same site in different lineages, this approach is expected to underestimate the number of mutation events, in particular for recurring mutations. Similarly to before, we classify three classes of sites with a variant allele: (i) those where the variant allele is present in any number of sequences; (ii) those where it is present in at least two sequences; (iii) those where it is present in at least five sequences.

Another issue that needs to be considered when estimating mutation rates is that different mutations can have different effects on the ability of the virus to replicate and transmit. In this manuscript, we mainly focus on synonymous mutations, as they are sufficiently abundant in our dataset to allow reliable estimates of mutation rates, and as they are expected to usually have a more limited effect on the viral fitness [26], therefore providing more limited biases in the inference of neutral mutation rates.

Later in the manuscript we describe a method to investigate possible fitness effects of synonymous mutations in SARS-CoV-2. This method is based on the comparison of ratios of estimated numbers of mutations with different numbers of descendants. The principle behind this method is that negative selection tends to decrease the frequency of new mutations, while positive selection tends to increase it. So, for example, if we want to compare the fitness of synonymous C→U mutations versus A→C ones, we can compare the ratio of high-frequency vs low-frequency synonymous C→U mutations, to the same ratio for synonymous A→C mutations. If the former is significantly higher, we take this as evidence that synonymous C→U mutations have typically higher fitness than A→C ones.

More details on our approaches are given in the Results and Material and Methods sections below.

## 3 Results

### 3.1 Neutral mutation rates in SARS-CoV-2

Here, we want to estimate the underlying mutability of different nucleotides, in a way that is as unbiased as possible with regards to how these mutations might affect the ability of new mutations to replicate and spread. To do this, firstly, we mostly focus on synonymous mutations, which are expected to affect, on average, the fitness of the virus much less than nonsynonymous mutations [26]. Secondly, we only consider new SARS-CoV-2 mutations observed within the human population, and ignore long-term divergence (between-species substitutions) which are expected to be more affected by selective forces [27]. While we cannot exclude that selection still affects some of the patterns observed below, for example making some types of synonymous mutations more lethal for the virus than others, we tried to reduce these biases as much as possible in the following.

First, to put our results in perspective of previous studies, we looked at numbers of sites with alternative alleles. When looking at patterns across the whole genome, it appears that all transitions (C→U, U→C, A→G and G→A) are quite common, as well as G→U mutations; note however that A and U bases have more opportunities to mutate as they are more common in the genome (Figure 1A). If we focus only on possible synonymous mutations (which we expect to be less likely affected by strong selection), we see C→U and G→U synonymous alternative variants at the vast majority of sites at which such variants could have been observed (Figure 1B). This means that sites at which synonymous C→U and G→U mutations might have occurred are possibly saturated with such mutations. In fact, using our phylogenetic approach to estimate numbers of mutation events of each type at each genome position, we clearly see that C→U and G→U are the most frequent mutations (Figure 1C), despite having fewer opportunities to occur due to GC content being lower than AT content. Focusing again on synonymous mutations, we can see that, while we don’t infer more overall G→U mutations than U C or A G ones, given the very low number of sites at which synonymous G→U mutations are possible, G→U and C→U seem the synonymous mutations that occur proportionally more often (Figure 1D). We also observe similar patterns for non-coding, non-synonymous and 4-fold degenerate sites (Figure S1). These observations suggest that the G→U and C→U underlying neutral mutation rates are considerably higher than all others.

**Figure 1:**
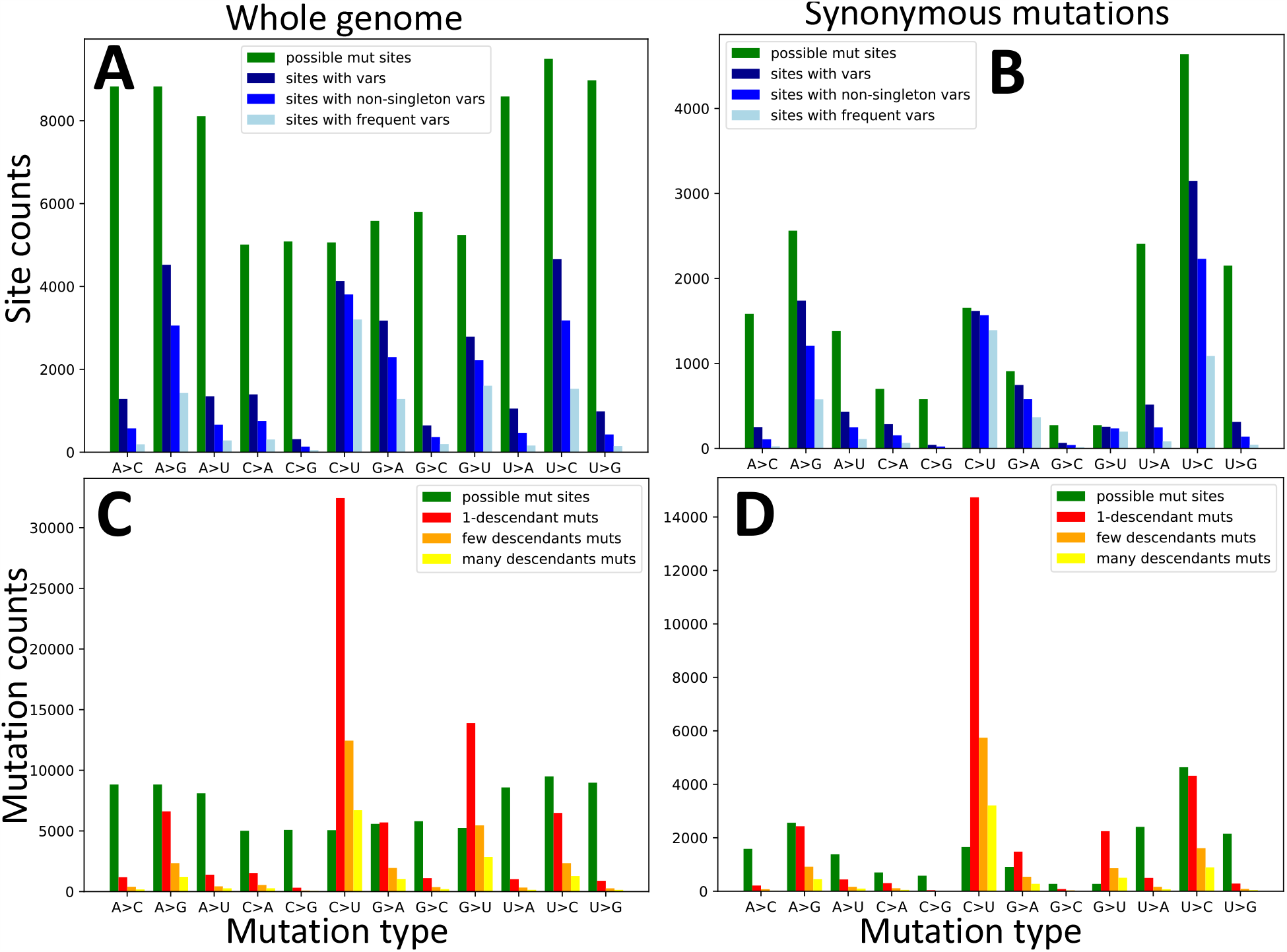
Numbers of possible mutations, observed mutations, and sites with alternative alleles. On the X axes are the 12 distinct types of mutation events, A→C, A→G, etc. In green we always show the number of genome positions at which the considered mutation type is possible. In **A,C** we consider all possible mutations, while in **B,D** we consider only synonymous mutations. In **A,B** we show, on the Y axis, the numbers of sites with alternative alleles in the alignment (blue color hues). In dark blue we show the number of all sites with alternative variants of the given type; in blue, we only show the number of such sites at which the alternative variant is present in at least 2 sequences; in light blue, only sites at which the considered alternative allele is present in at least 5 sequences. By definition, in plots **A-B** green bars are necessarily taller than all blue ones. In **C-D** we show, in red, orange and yellow, the numbers of mutation events inferred with parsimony on our phylogeny. In red we show the number of mutation events of the considered type with exactly one descendant; in orange the number of these mutations with at least 2 but less than 5 descendants; in yellow, those with at least 5 descendants. Mutation possibilities (green) can be fewer than inferred mutations events (red, orange and yellow in plots **C-D**) for certain types of mutations since the same mutation event can be inferred multiple times at the same site in different parts of the phylogenetic tree.

Our results further suggest that inferring mutation rates from counting the number of sites with variants (the counts in Figure 1A-B, and as in e.g. [5, 21]) can lead to underestimating the G→U and C→U synonymous mutation rates. This is because sites at which such mutations are possible are probably often saturated (multiple mutation events of exactly the same type have occurred along the phylogeny). However, saturation in previous studies was probably not as extreme as here since we investigate a considerably larger number of genomes. Another difference of our approach from most previous studies (but similar to [5]), is that we aim to disentangle the contribution of selection acting on the amino acid sequence from the underlying mutation rates, and to do this we separate synonymous and nonsynonymous mutations. This has a considerable further effect on the inference of the G→U mutation rates, because there are only a few sites at which a G→U mutation is synonymous, and so most possible G→U mutations are probably under significant purifying selection; skews in the genetic code [28, 29] are probably an important factor here. Of further impact, G is less frequent in the SARS-CoV-2 genome than U, in particular at 4-fold degenerate sites (sites where any 1-base mutation is synonymous) with a frequency of 6.5%, compared to frequencies of A 28.9%, C 11.4%, and U 50.9%; this causes a further under-representation of observed G→U mutations relative to the underlying G→U mutation rate.

A confirmation that many synonymous G sites are saturated with mutations can be seen in Figures 2 and S2. C→U and G→U mutations are by far the most homoplasic (the same mutation occurs more than once along the phylogeny), especially at synonymous and non-coding positions. This is unlikely to be the result of positive selection favouring these mutations, or of phylogenetic tree inference or parsimony mutation inference errors (since this seems to affect only C→U and G→U mutations) and seems instead the result of underlying relatively high neutral C→U and G→U mutation rates.

**Figure 2:**
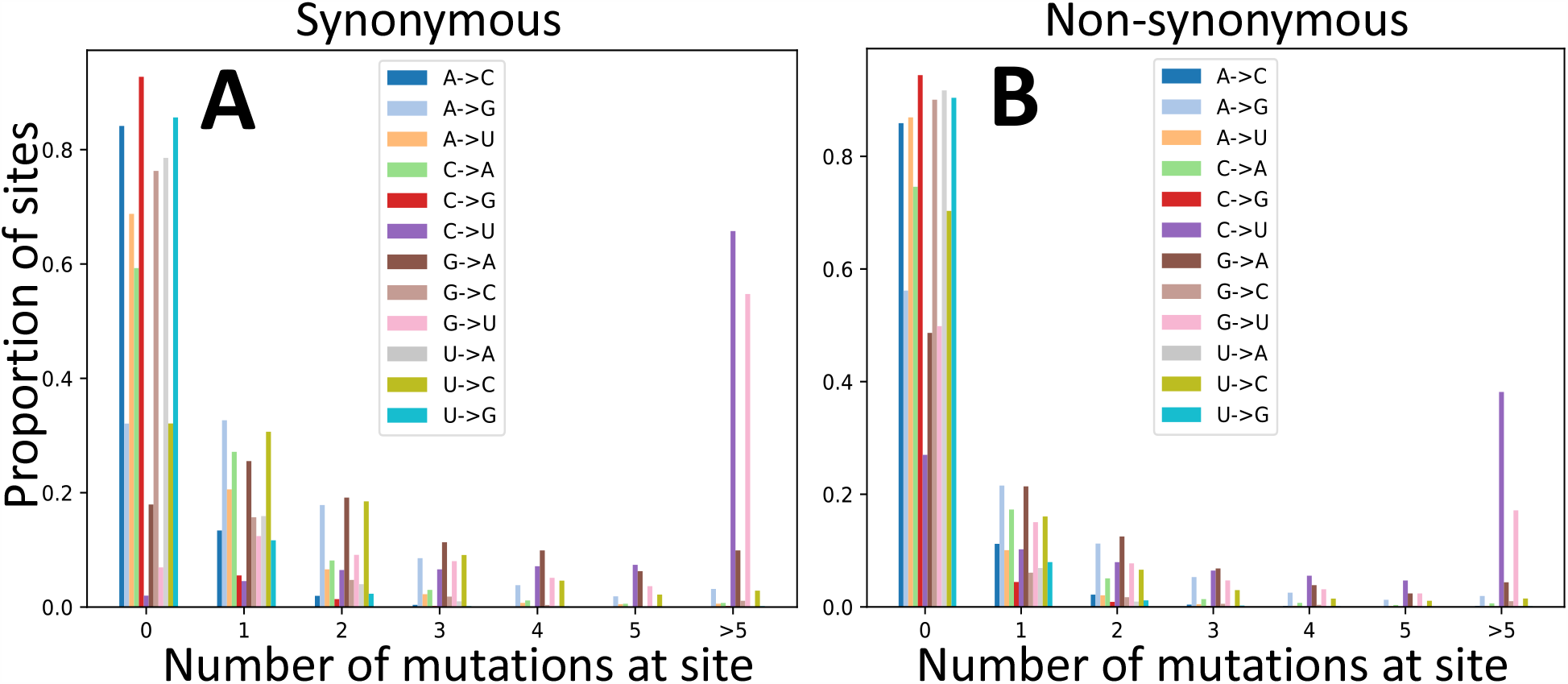
Re-occurrence of mutation events at the same sites. Proportion of sites (Y axis) where a given mutation (color, see legends) appears a certain number of times (X axis) along the phylogeny. **A** synonymous sites; **B** non-synonymous sites.

We also mostly find C→U or G→U mutations among the most homoplasic mutations of the genome: 55 out of 59 total mutations that occur more than 60 times at the same position are C→U or G→U. Here we ignore sites that are so homoplasic as to probably cause issues in phylogenetic inference [23, 22]; these sites were masked before any analysis here, and are namely the G→U mutation at position 11083, and the C→U mutations at positions 16887 and 21575. These three sites appear as the most mutable in the genome. However, the next 3 most homoplasic mutations of the genome that we identify here are neither C→U nor G→U mutations. The most recurrent one is the A→G nonsynonymous (K→R) mutation at position 10323 which we inferred to have occurred 138 times, with a total of 1187 descendants, found in context CTTAAGCTTA<u>A</u>GGTTGATACA. The second is a A→G nonsynonymous (K→R) mutation at position 21137, which occurred 130 times with a total of 472 descendants, found in context ATACAACAAA<u>A</u>GCTAGCTCTT. The third is a T→C synonymous mutation at position 27384 which occurred 119 times, with a total of 808 descendants, found in context TGGAGATTGA<u>T</u>TAAACGAACA. We suspect that these 3 mutations are the results of frequent ADAR activity, considering also their context. For example, the first two are A→G mutations with a G downstream and an A upstream (see [30, 31]). The third site is probably affected by ADAR acting on the negative strand. The fourth highly homoplasic mutation we found that is not G→U or C→U is the 11th most common in the genome, a G→A nonsynonymous (G→S) mutation at position 1820, with a total of 409 descendants and in context AGCTAAAAAA<u>G</u>GTGCCTGGAA.

One of the principal aims of our work is to estimate mutation rates while taking into account the effect of selection acting on the amino acid sequence, and trying to account as much as possible for other issues such as homoplasic mutations. To do this, firstly, we focus on synonymous mutations, which are expected to be less subject to selective constraints than nonsynonymous ones, while being much more abundant than non-coding mutations. Secondly, we use inferred counts of mutation events (using parsimony inference along our phylogenetic tree, see Methods); this accounts for the saturation of mutation events at more mutable positions, as discussed before. Thirdly, we separate mutation counts according to the number of observed descendants of each mutation - this allows us to have independent estimates, and to have estimates that do not rely on inferred mutation events with one descendant, which might be enriched in sequence errors or RNA degradation (in the case errors RNA degradation would be present in our alignment). Fourthly, we normalize mutation counts by the number of mutation possibilities, so to account for the fact that certain mutation types (for example synonymous G→U mutations) are possible at fewer sites than other mutation types. For this last point, we divide the number of inferred mutations of a certain type (e.g., red, orange and yellow bars in Figure 1D) by the number of sites at which such mutations are possible (e.g., green bars in Figure 1D), always considering the reference genome as the mutational ancestral background. We find that the G→U (transversion) mutation rate is similar to C→U (transition) (Figures 3 and S3), and they are both considerably higher than all other rates (about 4 times higher than the next highest rate, G→A, see Table 1). We confirm that all other transitions, U→C, G→A and A→G, have lower rates, but higher than all remaining transversions. In particular, the mutational process seems highly asymmetrical and strand-asymmetrical, with mutation rate G→U being 82.0 times higher than U→G and 21.1 times higher than C→A (Table 1). This strand-asymmetry is likely the result of ROS and APOBEC activity on single-strand RNA [14, 15].

**Table 1:** Mutation rates estimated from 4-fold degenerate sites. Only mutation events with more than 1 descendant have been considered here. Rates are been normalized as typically done in phylogenetics, the normalizing constant being the sum of the rates multiplied by the frequency of the ancestral allele: ∑_*a,b*_ *q*_*a,b*_*π*(*a*), where *a, b* are nucleotides, *q*_*a,b*_ is the original unnormalized rate from allele *a* to *b*, and *π*(*a*) is the frequency of *a* at 4-fold degenerate sites.

|  | to A | to C | to G | to U |
| --- | --- | --- | --- | --- |
| from A |  | 0.039 | 0.310 | 0.123 |
| from C | 0.140 |  | 0.022 | 3.028 |
| from G | 0.747 | 0.113 |  | 2.953 |
| from U | 0.056 | 0.261 | 0.036 |  |

**Figure 3:**
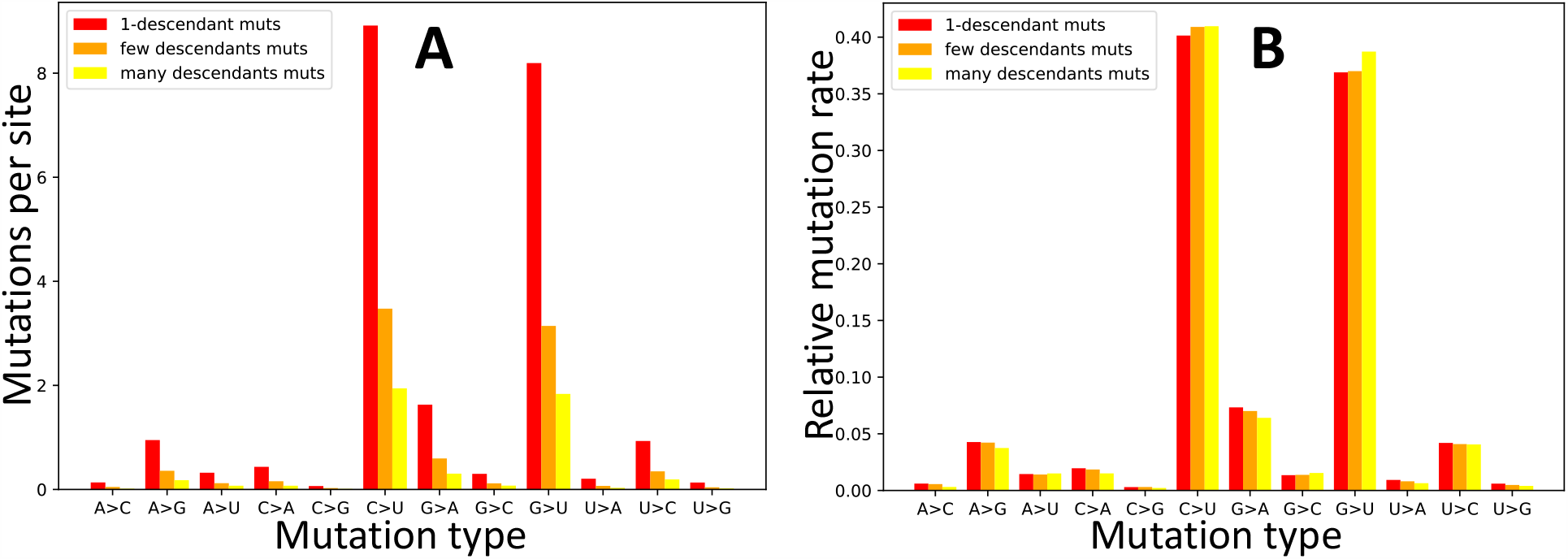
Estimated synonymous mutation rates in SARS-CoV-2. To estimate synonymous mutation rates in SARS-CoV-2 we used the counts of inferred synonymous mutation events (see Fig 1D) normalized by the numbers of reference genome sites at which such mutations might have occurred. On the X axis are the 12 distinct types of mutation events, A→C, A→G, etc. In red, orange and yellow we show respectively rates obtained from counts of mutation events with 1 descendant, more than 1 but less than 5 descendant, and 5 or more descendants. **A** Mutation rates represented as average numbers of mutation events inferred per site at which such mutation type is possible. **B** Relative mutation rates (the sum of all bars of one specific color is 1.0).

Our results differ remarkably from those studies that either estimated mutation rates by comparing numbers of sites with alternative alleles, did not divide these counts by the numbers of opportunities for such mutations, and/or did not account for the biases in the genetic code by separating synonymous and nonsynonymous mutations (e.g. [20, 32]). Our results are instead more consistent with those of studies that did take some of these steps and highlighted similarly high C→U and G→U mutation rates in SARS-CoV-2 (e.g. [5]). While elevated C→U mutation rate in SARS-CoV-2 has been frequently observed and usually attributed to the effects of APOBEC activity, the elevated G→U has been discussed much less, but it has been usually suggested to be the result of the activity of reactive oxygen species (ROS), see [5, 15]. The number of descendants of mutation events seems to have little impact on our inferred relative mutation rates (Figure 3B); this suggests that phenomena like RNA degradation or sequence errors, which are expected to overwhelmingly result in inferred mutation events with one descendant, do not considerably affect our mutation rate estimates.

### 3.2 Context dependencies

One of the observations that suggests APOBEC activity being the leading cause of the elevated C→U mutation rate in SARS-CoV-2 is that nucleotide context seems to affect the C→U mutation rate in a way that is consistent with the action of some of the human APOBECs [15]. Here we study the effect of neighbouring base context on C→U mutability. We divide the set of possible and observed C→U mutations in classes based on the nucleotide context (5’ to 3’ preceding and following base). Even though we focus only on C→U mutations, it is still important to separate synonymous and nonsynonymous mutations, since different contexts can lead to synonymous or nonsynonymous C→U mutations with different proportions. Additionally, different contexts can be present with different frequencies in the SARS-CoV-2 genome. Focusing on synonymous mutations, and using the normalization procedure outlined above, we find that U<u>C</u>G→U<u>U</u>G appears to be the most mutable context (Figures 4A and S4). Note that this pattern is not observed if one considers unnormalized mutation counts, particularly those from nonsynonymous mutations (Figure S5).

**Figure 4:**
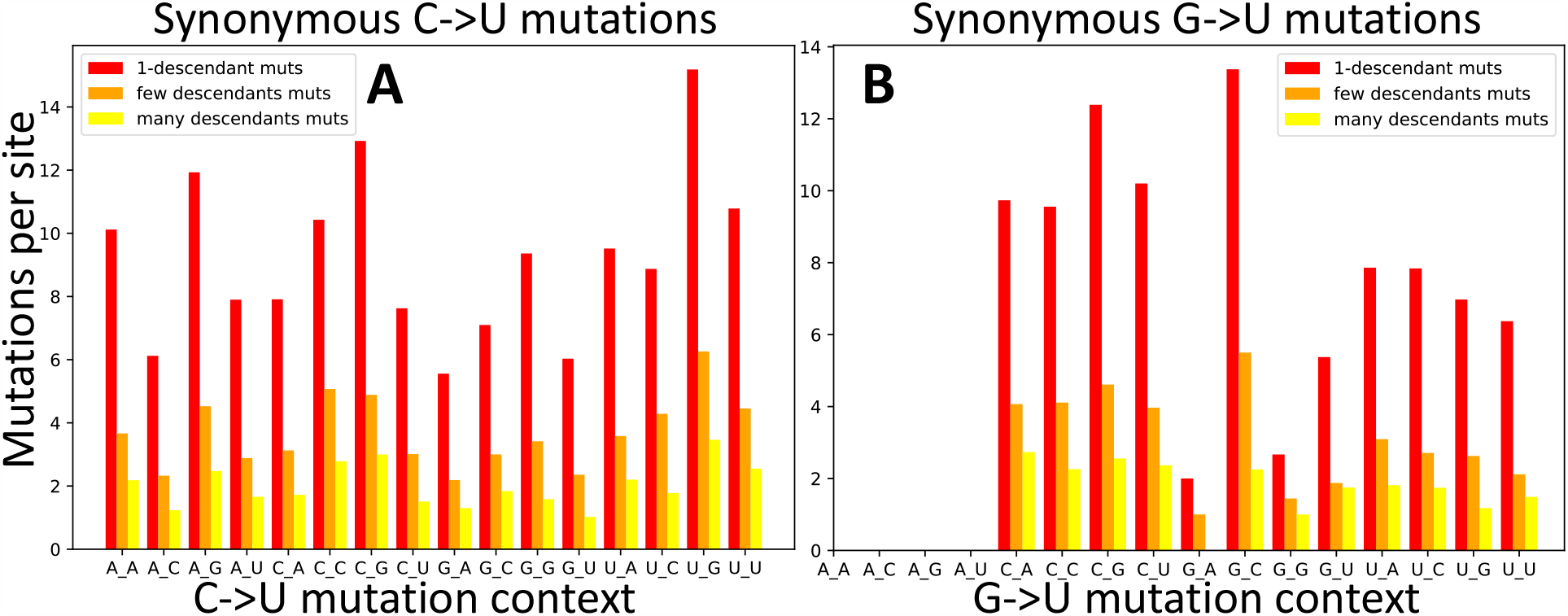
C→U and G→U synonymous mutation rates in different base contexts. Here mutation rate are calculated as in Figure 3A. **A** C→U mutations. **B** G→U mutations. The X axis shows the context of the considered mutation (for example, in **A**, A_G represents the trinucleotide A<u>C</u>G and its synonymous mutation rate into trinucleotide A<u>U</u>G). Colors are as in Figure 3.

This confirms previous results which suggested an elevated U<u>C</u>G→U<u>U</u>G mutation rate [32, 16, 5], attributed to either the context-specificity of APOBEC mutational targets, or to selection against CpG dinucleotides [15]. We discuss selection in the next section. Here, we note that while we found a signal of context affecting mutation rates in SARS-CoV-2, and while at least part of this observation is consistent with some of the known sequence targets of APOBEC, overall, C→U mutations occurring in U<u>C</u>G or more generally U<u>C</u>→<u>U</u>U context represent only part (and a very small part for U<u>C</u>G) of all C→U mutations in SARS-CoV-2 (Figure S1) and as such they don’t explain by themselves the whole C→U relative hypermutability.

When we look at longer sequence context (previous 5 bases and following 5 bases) we see that U in the 2 bases preceding a C seems to have a C→U mutagenic effect, while G in the previous 2 bases and C in the following 2 bases seem to reduce the mutation rate (Figure S6).

Given the relatively high G→U mutation rate, we performed a similar analysis of context effects on G→U mutations. However, while having a high rate, G→U synonymous mutations are quite rare (Figure S7 for numbers in different contexts) and so estimates of context-dependent rates are expected to have substantial uncertainty. Additionally, some contexts are not possible for G→U synonymous mutations, for example AG→AU mutations are never synonymous (Figure S7). We don’t notice any one particularly elevated mutational G→U context, however, a G being preceded or followed by a C (C<u>G</u> and <u>G</u>C contexts) seems to have a higher G→U mutation rate (Figures 4B and S8).

### 3.3 Testing for selection at synonymous sites using mutation frequencies

Previous authors have discussed the effect of selection on synonymous mutations in SARS-CoV-2. For example, the elevated mutation rate in CpG context has been interpreted as a sign of selection against CpG content in SARS-CoV-2 in order to avoid zinc finger antiviral proteins (ZAP), see [15]. This is consistent with similar evidence found from depleted CpG content in other coronaviruses [33]. However, evidence from CpG content and substitution rates can be confounded by skewed mutation rates and mutational contexts like the ones due to APOBEC activity, which might be the leading cause of mutation in SARS-COV-2.

To overcome this limitation, and in order to disentangle the contribution of mutation and selection over the synonymous evolution of SARS-CoV-2, [5] compared equilibrium frequencies inferred from SARS-CoV-2 substitution rates to observed nucleotide and dinucleotide frequencies. Since selection tends to raise the frequency of favourable alleles in a population, and to decrease the frequency of deleterious alleles, if a certain nucleotide is advantageous over another at synonymous sites, for example, if C is advantageous over U, it is expected that relatively more U→C mutations will reach fixation (completely replace the ancestral allele in the population) than C→U mutations. This means that, in the long term, there would be fewer U nucleotides in the genome than expected based on the mutation rates estimated from observed genetic variation within the population. [5] observed that U nucleotides are less common at 4-fold degenerate sites in SARS-CoV-2 than expected from SARS-CoV-2 genetic variation within humans, and concluded that there is ongoing selection against U nucleotides in SARS-CoV-2. With a similar analysis, they also concluded that there is no ongoing selection against CpG content in SARS-CoV-2, despite opposite prior expectations (see [15, 33]).

[5] assumed that the evolutionary process is stationary, that is, that mutation rates and selective pressures did not change recently (for example, since the introduction of SARS-CoV-2 into humans). This assumption is highly debatable. Due to the recent host shift and to the fact that many mutations in SARS-CoV-2 seem to be the consequence of host immune system activity, a recent significant change in mutation rates and selective pressure in SARS-CoV-2 (associated with its introduction to humans) is likely. In fact, studies have suggested that the current G→U mutation rate of SARS-CoV-2 in humans, for example, is much higher than in its reservoir hosts [34, 35]. Specifically, [34] estimated that the G→U mutation rate in SARS-CoV-2 increased 9-fold with its introduction in humans, possibly due to difference in ROS activity between hosts.

We used the approach of [5] on our dataset to estimate mutation rates at 4-fold degenerate sites, and then from these estimated equilibrium nucleotide frequencies using the python package discreteMarkovChain v0.22 https://pypi.org/project/discreteMarkovChain/. Similarly to [5], we find that 4-fold degenerate site U content is considerably lower than the equilibrium U content (50.9% vs 65.5%). However, if we re-calculate the equilibrium nucleotide frequencies after decreasing the G→U mutation rate 9-fold in this analysis (to mimic the putative reservoir mutation rate estimated in [34]), the difference almost disappears (equilibrium U content 52.6%). If we repeat the analysis above using mutation rates as in Table 1 (inferred from mutation counts) instead of mutation rates inferred as in [5] (inferred from the numbers of sites with variant alleles, as in Figure 1B), we find an equilibrium U frequency of 77.4%, and, after decreasing G→U rate 9-fold, of 64.9%. These observations suggest that, indeed, changes in mutation rates can cause strong changes in equilibrium frequencies, and the inference of selection from the comparison of current and equilibrium nucleotide frequencies can be affected by changes in mutation rates.

To address this issue, we propose an alternative approach to test for selection acting on synonymous mutations. Since selection increases the frequency of favourable alleles and decreases the frequency of deleterious alleles within a population, we expect selection to change the relative proportion of alleles at different population frequencies. The McDonald-Kreitman test [27], for example, compares numbers of synonymous and nonsynonymous within-species polymorphisms and between-species polymorphisms; within this framework, purifying selection is expected to decrease the number of nonsynonymous substitutions more than the number of nonsynonymous polymorphisms.

However, the emergence of SARS-CoV-2 in humans is relatively recent, too recent for variants (and especially a statistically sufficient number of variants) to have reached fixation within the human population, making this type of test not applicable. Instead, we use an alternative version in which we compare low-frequency mutations against high-frequency ones. While our aim is to focus on comparing different types of synonymous mutations, we first apply this approach to compare synonymous and nonsynonymous mutations, so as to make the similarity to the McDonald-Kreitman test more apparent. Nonsynonymous mutations in SARS-CoV-2 appear significantly shifted towards lower frequencies (Figure 5), consistent with the expectation that purifying selection tends to decrease the frequencies of nonsynonymous mutations more than the frequencies of synonymous mutations.

**Figure 5:**
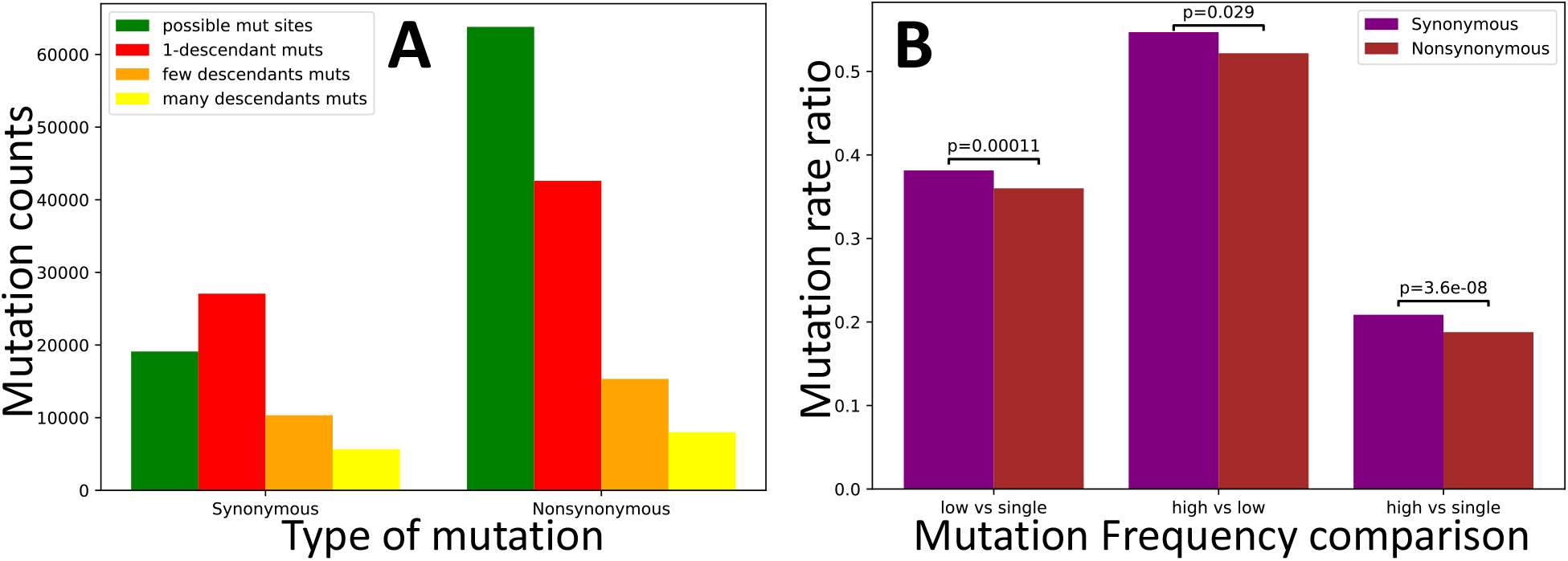
Evidence of selection affecting the population frequency of synonymous vs nonsynonymous mutations. Counts and rate ratios of SARS-CoV-2 synonymous and non-synonymous mutations at different frequencies in the human population. **A** counts of possible mutations (green), singleton mutations (red), low frequency (> 1 and 4≤ descendants) mutations (orange), and high frequency (> 4 descendants) mutations (yellow). **B** ratios of higher vs. lower frequency mutation rates. In the absence of selection, ratios should not be significantly different between the classes of synonymous and nonsynonymous mutations. Instead, we measure a significant deviation in each comparison, with nonsynonymous mutations being relatively depleted of high frequency mutations. We calculated p-values using the chi2_contingency function of the Scipy.stats package [36].

Next, we focused on the hypothesis that U variants at synonymous sites are on average mildly deleterious [5]. This time, not all of our comparisons are significant, but those that are, consistently suggest that selection favours U variants (Figure 6), not the opposite. As we discussed before, the difference with the estimate from [5] probably lies in the assumption of stationarity in SARS-CoV-2 evolution (although we also used a larger dataset and a different approach to estimate mutation rates).

**Figure 6:**
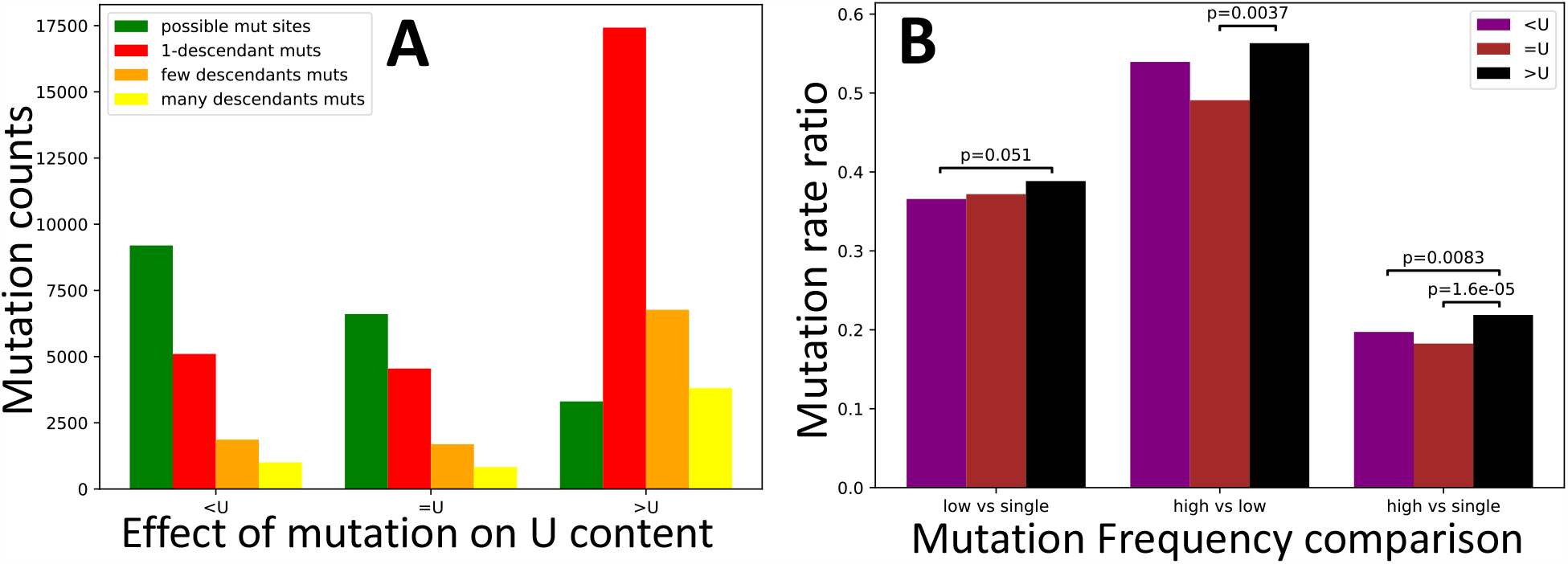
Test of selection affecting U content at synonymous sites. Values are the same as in Figure 5, but this time we focus on synonymous mutations that decrease U content (“*<*U”), increase it (“>U”), or leave it unaltered (“=U”). Only p-values below 0.1 are shown.

A possible explanation for the pattern we observe is that many mutations increasing U content are C→U or G→U, and therefore may sometimes reduce CpG content. Selection against CpG content has already been suggested in coronaviruses [33], and is expected due to ZAP activity [15]. However, using our approach the evidence in support of this hypothesis is not significant (Figure S9). Similarly, when investigating possible selection affecting GC content, only one of the comparisons is significant (Figure S10).

## 4 Discussion

In this study, we investigated mutation rates and selection at synonymous sites in SARS-CoV-2. We used a different methodology than previous studies, so as to exploit more than 147, 000 SARS-CoV-2 genome sequences while trying to avoid inaccuracies due to mutation saturation. Our approach also accounts for the fact that different types of mutations can have, on average, different fitness due to their impact on the amino acid sequence and due to skews in the genetic code. Also, we did not assume equilibrium in SARS-CoV-2 genome evolution, which is unlikely given the recent shift of host the virus underwent.

We found that two mutation rates, C→U and G→U, are similar to each other and much higher than all others mutation rates, leading to extremely frequent homoplasies. This also means that the SARS-CoV-2 mutation process is very far from being symmetric or strand-symmetric; it is also considerably far from equilibrium. We also found some sites with extremely high non-C→U non-G→U mutation rates that are consistent with targeted ADAR activity.

A consequence of these findings is that popular and efficient phylogenetic substitution models such as JC69 [37], HKY85 [38], or GY94 [39] might be inappropriate with SARS-CoV-2 data and might cause biases in the inference of phylogenetic trees [23], positive selection [9] and recombination [12] due to the neutral high recurrence of certain mutations. It is therefore probably important to adopt phylogenetic substitution model that can account for elevated C→U and G→U rates, such as the non-reversible UNREST model [40], which however we found to usually cause numerical instability in current phylogenetic packages that allow it [41, 42]. In the future, if we want to alleviate these biases, it will be important to implement such more general substitution models more broadly and in a more numerically stable way.

When investigating selection at synonymous sites possibly affecting CpG or GC content, we found mostly non-significant patterns. We also investigated the possibility of selection against U content in SARS-CoV-2 [5], but found significant evidence in the opposite direction. While further analyses will be needed to establish with confidence what kind of selection acts non SARS-CoV-2 synonymous mutations, we suggest that inference based on assumption of genome equilibrium can be biased by changes in mutation rates and selective pressures associated with host shift. While we tried to account for possible biases as much as possible, our methods still have some limitations. First of all, our inference of mutation events is based on a prior phylogenetic inference, but tree inference from SARS-CoV-2 data is typically not very reliable, in part due to the low genetic diversity among sequences, but also due to homoplasic mutations [43, 23]. As mentioned above, our phylogenetic inference might also have been negatively affected by the choice of substitution models; however, currently, more realistic models like UNREST are either not implemented or are numerically unstable in sufficiently efficient phylogenetic packages such as [41, 44, 42]. In this study we tried not to rely excessively on individual inferences of mutation events, but rather focused on general patterns averaged over many sites and clades, which we think should provide robust inference despite the fact the inference of individual mutation events might not be reliable. However, a potential bias that might affect our result derives from the fact that some sites are very homoplasic, and our phylogenetic inference might lead to an over-parsimonious inference of their mutational history. This, in turn, might lead us to underestimate their mutation rate and overestimate their number of descendant tips per mutation events. In the future, a Bayesian phylogenetic approach might be useful to assess and possibly resolve this issue and assess its impact on our inference of selective pressure; however so far Bayesian phylogenetic inference has proved prohibitive with datasets of this size.

To further investigate and disentangle selective and mutational forces in SARS-CoV-2, it would be very promising to combine an analysis of between-patients and within-patient SARS-CoV-2 genetic variation. In a similar way as selection is expected to decrease the frequency of deleterious SARS-CoV-2 alleles at the human population level, the same is true at the within-host levels, as selection is expected to act on within-host deleterious mutations and often prevent them from reaching high frequency and transmit further on. While within-patient genetic diversity data can indeed be very informative of the SARS-CoV-2 evolutionary patterns [13, 14, 45], it is important to consider that such data is also more prone to sequencing, read processing, and RNA degradation issues. These issues cause some errors in consensus sequences [23, 22] but they are expected to be even more problematic at the level of detected within-host variation. Indeed, so far there is reason to be cautious when interpreting these data [22], especially with specific datasets (see https://virological.org/t/gained-stops-in-data-from-the-peter-doherty-institute-for-infection-and-immunity/486). Given reliable data regarding evolution at different levels (e.g. within a patient, between patients, and between hosts), it would be interesting to combine these sources of information to improve estimates of selective pressures (see e.g. [46, 47]).

Our methodology to detect selection could also be improved by using the full site frequency spectrum of mutations, instead of categorizing mutations into frequency classes (see e.g. [48, 49]). However, it is important to consider that complex epidemiological dynamics and sampling biases in SARS-CoV-2 mean that it is hard to interpret the shape of any individual site frequency spectrum in terms of the effects of mutation and selection. We infer mutation and selection patterns by comparing properties of site frequency spectra associated with different mutation types. This approach should be robust to the effects of variable population dynamics and sampling biases, since these forces are expected to affect in the same way site frequency spectra associated with different mutation types.

In the future, we hope to extend our current approach to also study insertions and deletions (indels): their frequency, recurrence and possible fitness effects (see e.g. [50]). However, we expect that incorporating indels in our approach will be challenging since standard phylogenetic approaches rarely model indels.

## 5 Materials and Methods

### 5.1 Data collection and phylogenetic inference

Full details and reproducible code for the construction of the global tree of SARS-CoV-2 samples are available in the 13/11/20 release of [51]. To summarise, this code creates a global phylogeny of all available samples from the GISAID data repository as follows.

First, all sequences marked as ‘complete’ and ‘high coverage’ submitted up to 13/11/20 were downloaded from GISAID. Sequences with known issues from previous analyses were then removed from this database (details are in the excluded_sequences.tsv file at the above DOI).

Second, a global alignment was created by aligning every sequence individually to the NC_045512.2 accession from NCBI, using MAFFT v 7.471 [52], faSplit (http://hgdownload.soe.ucsc.edu/admin/exe/), faSomeRecords (https://github.com/ENCODE-DCC/kentUtils), and GNU parallel [53]. This approach aligns each sequence individually to the reference, then joins them into a global alignment by ignoring insertions relative to the reference.

Third, sites that are likely to be dominated by sequencing error [23] are masked from the alignment using faSplit, seqmagick (https://seqmagick.readthedocs.io/en/latest/), and GNU parallel, sequences shorter than 28KB or with more than 1000 ambiguities are removed from the alignment using esl-alimanip (hmmer.org), and subsequently sites that are > 50% gaps are removed (after converting N’s to gaps) using esl-alimask.

Fourth, the global phylogeny was estimated using IQ-TREE 2 [41] FastTree 2 [44] (v2.1.10 compiled with double precision) in two stages. First, new sequences added to GISAID between 11/11/20 and 13/11/20 were added to the phylogeny inferred on 11/11/20 using Maximum Parsimony placement in IQ-TREE 2. This produces a starting tree of all sequences available on 13/11/20. Second, the starting tree was optimised using FastTree 2 with 2 rounds of Subtree Pruning and Regrafting (SPR) using moves of length 1000 under a minimum evolution optimisation regime, and the tree was then further optimised using multiple rounds of Maximum Likelihood Nearest Neighbour Interchange (NNI) moves until no further improvement to the tree could be achieved using NNI. The resulting tree was rooted with our reference (NC_045512.2 / MN908947.3 / Wuhan/Hu-1) sequence using nw_reroot [54].

From the resulting tree, we removed sequences on very long branches using TreeShrink [55] using the default, *q <*= 0.05 threshold, to identify such branches. These sequences are likely to be either of poor quality and/or poorly aligned, so rather unreliable to interpret in a phylogeny with such limited variation. The final tree and its related alignment contains 147,137 SARS-CoV-2 genomes.

### 5.2 Estimation of mutation rates

To separate mutation events into different categories, each position of the reference genome (NC_045512.2 / MN908947.3 / Wuhan/Hu-1, see https://www.ncbi.nlm.nih.gov/nuccore/MN908947) was classified as coding or non-coding. Start and Stop codons were not considered in the following analysis. The first and last 100bp of the genome, in addition to sites marked as problematic in https://github.com/W-L/ProblematicSites_SARS-CoV2/blob/master/problematic_sites_sarsCov2.vcf [22, 23], were also not considered here. We counted “possibilities” of mutations based on the reference genome: for example a non-coding C allele in the reference genome represents 3 possibilities for non-coding mutations (C→A, C→G and C→U). For coding sites, we split synonymous and non-synonymous mutation possibilities into two separate counts. Similarly, we also used the reference genome to define the number of possible mutations within each genetic context. Sites that were masked in the alignment were still used here to define the genetic context of possible mutations at neighbouring non-masked sites.

We then inferred a mutational history with parsimony over our global maximum likelihood tree, as described in [23]. The software for doing this analysis is available from https://github.com/yatisht/strain_phylogenetics. Given this inferred mutational history, for each site, we counted all the mutation events inferred from the reference allele to any alternative allele, and we recorded the number of descendant tips of each mutation event, excluding those that are descendants of further downstream mutations at the same site. Note that we do not count mutation events that modify a non-reference allele. For example, if at a U position of the genome we have a U→C mutation, and in one of the descendants of this mutation event we have a C→A mutation at the exact same position, then we count the first U→C mutation but not the second C→A; this is done to aid the normalization of mutation counts, see below. At each site and for each mutation type, we kept counts of three different classes of mutations: (i) those with only one descendant tip (“singletons”); (ii) those with ≥2 and ≤ 4 descendants (“low frequency”); and (iii) those with > 4 descendants (“high frequency”). Number of different mutation types and mutation possibilities observed across the SARS-CoV-2 genome are given in Figures 1 and S1. Note that the number of observed mutations is often higher than the number of mutation possibilities. For example, a reference 4-fold degenerate site with an reference A allele counts as one A→C, one A→G and one A→U synonymous mutation possibility. However, an A→C mutation event can occur multiple times along the phylogeny at this site, possibly resulting in multiple A→C synonymous mutations observed. When classifying mutation events into different categories, we assume that mutations happen in the genetic background of the reference genome. This might be inaccurate in some cases, but given the overall low level of divergence (all reliable sequences so far being diverged less than 50 substitutions from the reference genome) the effect of this approximation should be very limited.

The low mutation rate observed in SARS-CoV-2 makes parsimony an efficient and reliable approach to infer mutational histories [56]. However, phylogenetic inference from large SARS-CoV-2 datasets is difficult due to elevated computational demand and phylogenetic uncertainty [43], and we cannot exclude the presence of errors in our phylogenetic tree, and therefore in our mutational history.

As mentioned earlier, we also consider an alternative approach to estimating mutation rates, based on counting the number of alignment columns at which alternative alleles are observed. We classify three classes of variant allele: (i) those present in any number of sequences; (ii) those present in at least two sequences; (iii) those present in at least five sequences. Numbers of different types of variable sites found in our alignment are given in Figures 1 and S1.

For each class of mutations (synonymous, nonsynonymous, or noncoding; from any nucleotide to any other nucleotide; within any sequence context) we estimate an approximate mutation rate by dividing its total mutation count by its possibilities count. These rates represent the average number of mutation events of a certain type expected per given site along the considered phylogeny.

## 6 Code Availability

The code and data used for this project (except for the sequences which fall under the restrictions of the GISAID terms of use) are available at https://bitbucket.org/nicolademaio/mut-sel_sc2/.

## 7 Acknowledgments

NG, CW and NDM were supported by the European Molecular Biology Laboratory (EMBL). R.C.-D. was supported by R35GM128932 and by an Alfred P. Sloan foundation fellowship. R.L. is funded by Australian Research Council grant DP200103151, and by a Chan-Zuckerberg Initiative grant.

We are very grateful to GISAID and all the groups who shared their sequencing data. A full list of acknowledgments is available from https://github.com/roblanf/sarscov2phylo/tree/master/acknowledgements.

## 8 Supplement

### 8.1 Supplementary Figures

**Figure S1:**
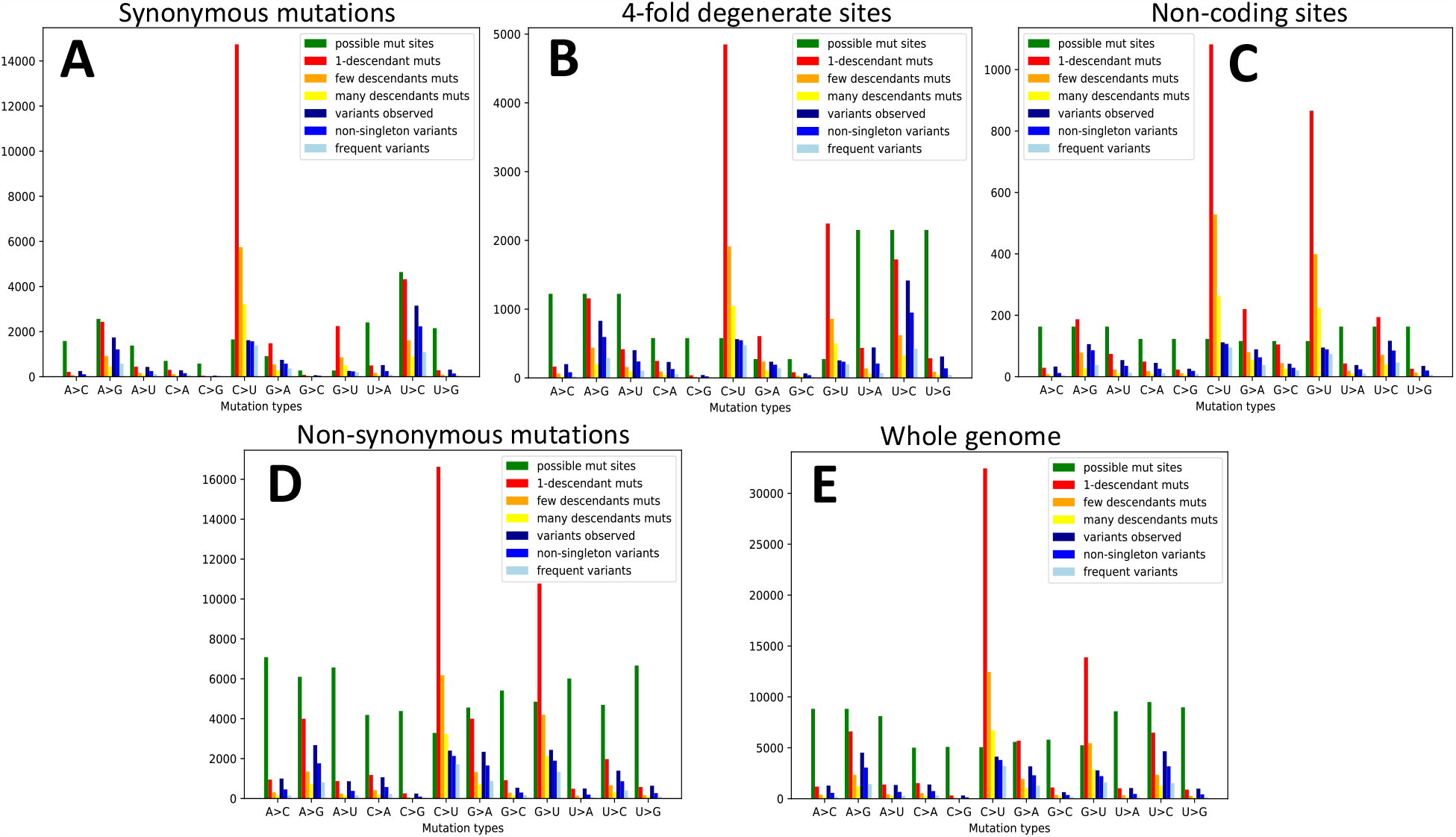
Numbers of possible mutations, observed mutations, and variable sites in the SARS-CoV-2 genome. Counts of mutation events for each site class: **A** synonymous sites, **B** 4-fold degenerate sites, **C** non-coding sites, **D** non-synonymous sites, **E** all sites. On the X axis are the 12 distinct types of mutation events, A→C, A→G, etc. In green we show the number of reference sites at which a mutation might have occurred. In red, orange and yellow we show respectively the number of observed mutations with 1 descendant, more than 1 but less than 5 descendants, and more than 4 descendants. In dark blue, blue, and light blue, we show respectively the number of sites with > 0, > 1, and > 4 variants of the given type.

**Figure S2:**
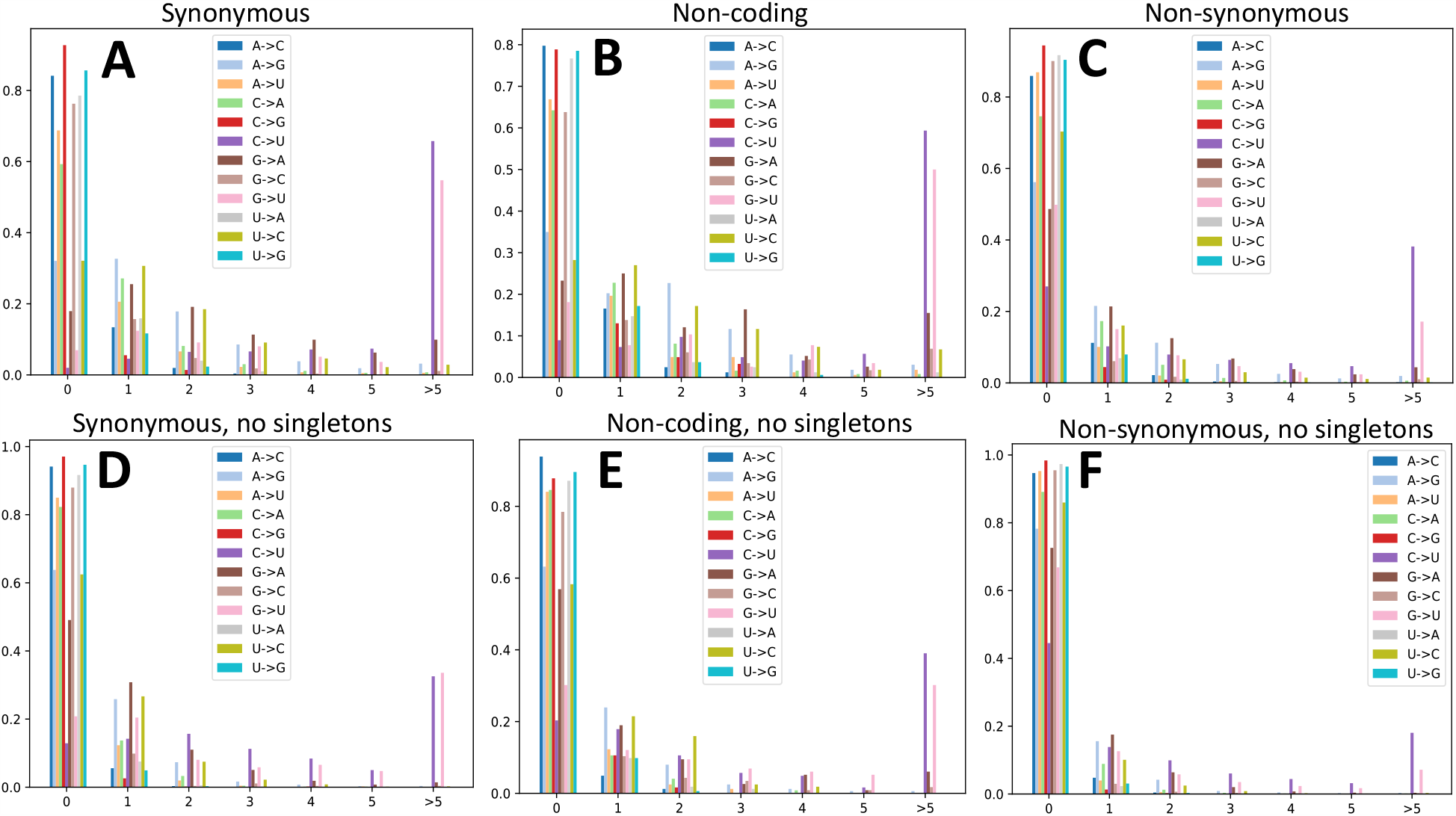
Re-occurrence of mutation events at the same sites. Here we show the proportion of sites (Y axis) where a given mutation (color, see legends) appears a certain number of times (X axis) along the phylogeny. **A** synonymous sites; **B** non-coding sites; **C** non-synonymous sites; **D** synonymous sites, but counting only mutation events with more than 1 descendant; **E** non-coding sites, only mutations with more than 1 descendant; **F** non-synonymous sites, only mutations with more than one descendant.

**Figure S3:**
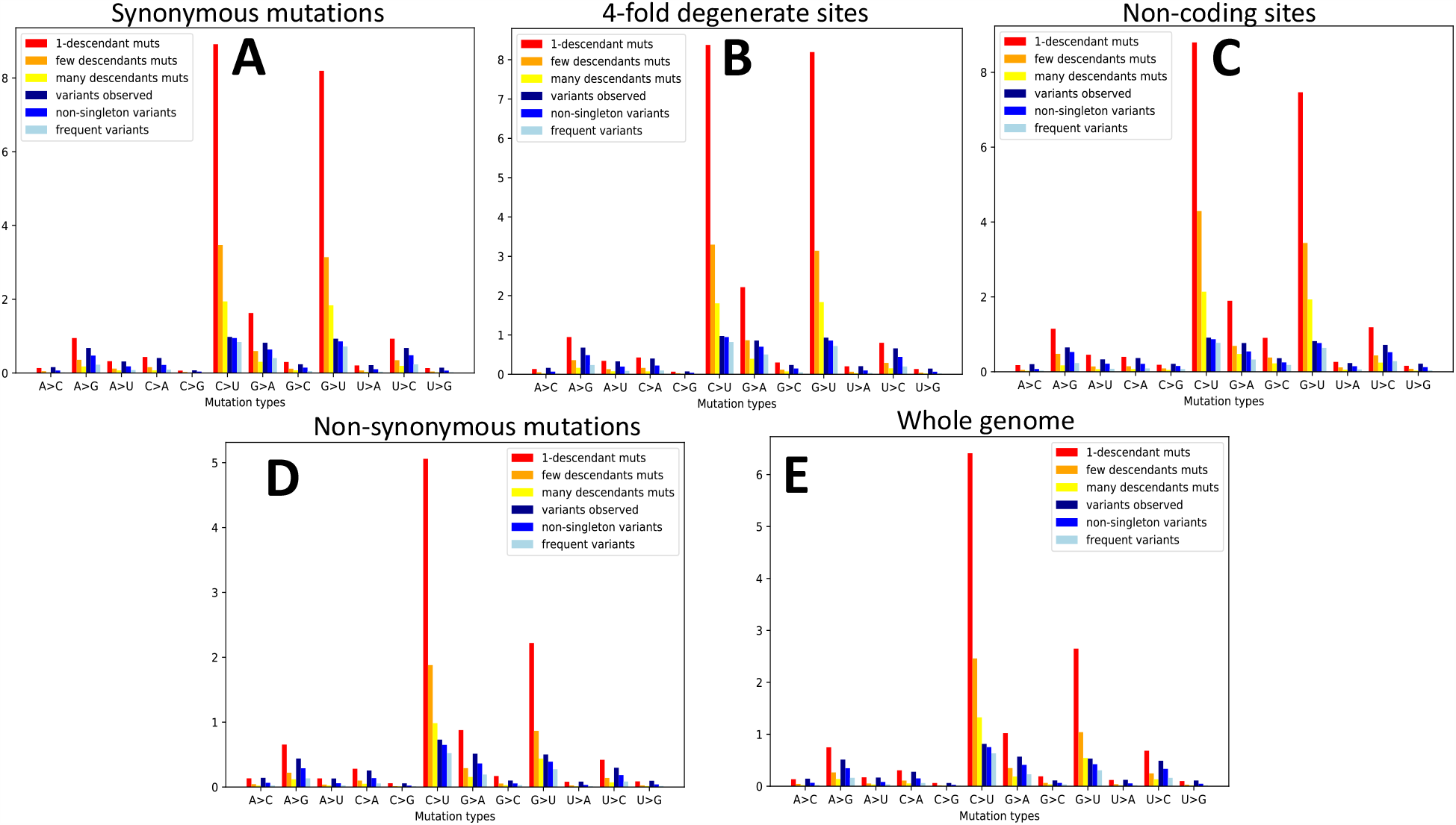
Mutation rates estimated from mutation counts and variable sites counts. On the X axis are the 12 distinct types of mutation events, A→C, A→G, etc. On the Y axis are the inferred mutation rates for **A** synonymous sites, **B** 4-fold degenerate sites, **C** non-coding sites, **D** non-synonymous sites, **E** all sites. In red, orange and yellow we show respectively the mutation rates inferred from the numbers of observed mutations with 1 descendant, more than 1 but less than 5 descendant, and more than 4 descendant (and dividing each count by the number of reference sites where such mutations might have happened). In dark blue, blue, and light blue, we show respectively the mutation rates inferred from the numbers of sites with > 0, > 1, and > 4 variants of the given type.

**Figure S4:**
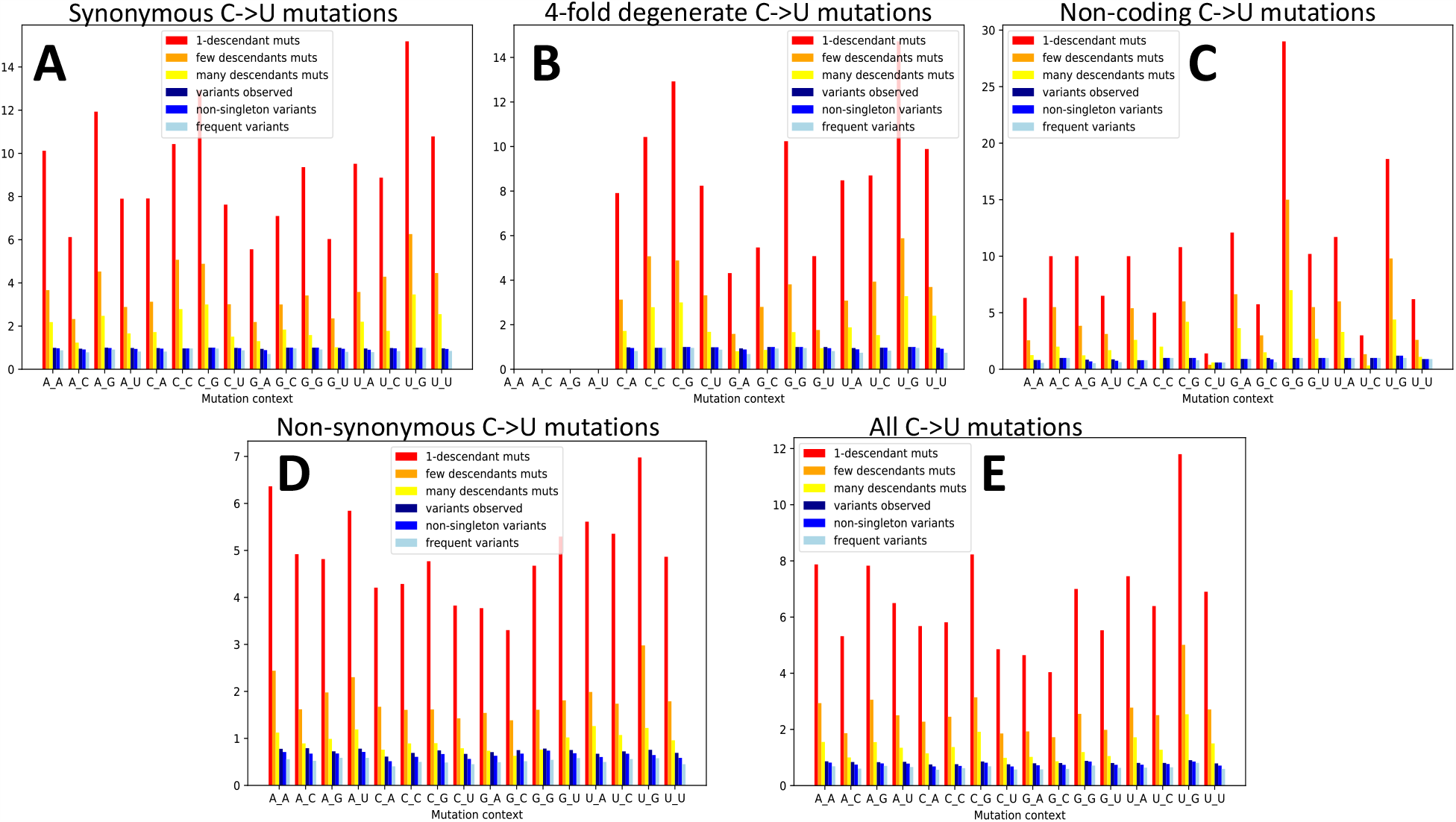
C→U mutation rates in different base contexts. C→U mutation rate depending on the previous and next base (5’ and 3’ base neighbours, shown on the X axis). A_G represents, for example, the trinucleotide ACG and its mutation rate into trinucleotide AUG. Colors are as in legend Figure 3. **A** synonymous sites, **B** 4-fold degenerate sites, **C** non-coding sites, **D** non-synonymous sites, **E** all sites.

**Figure S5:**
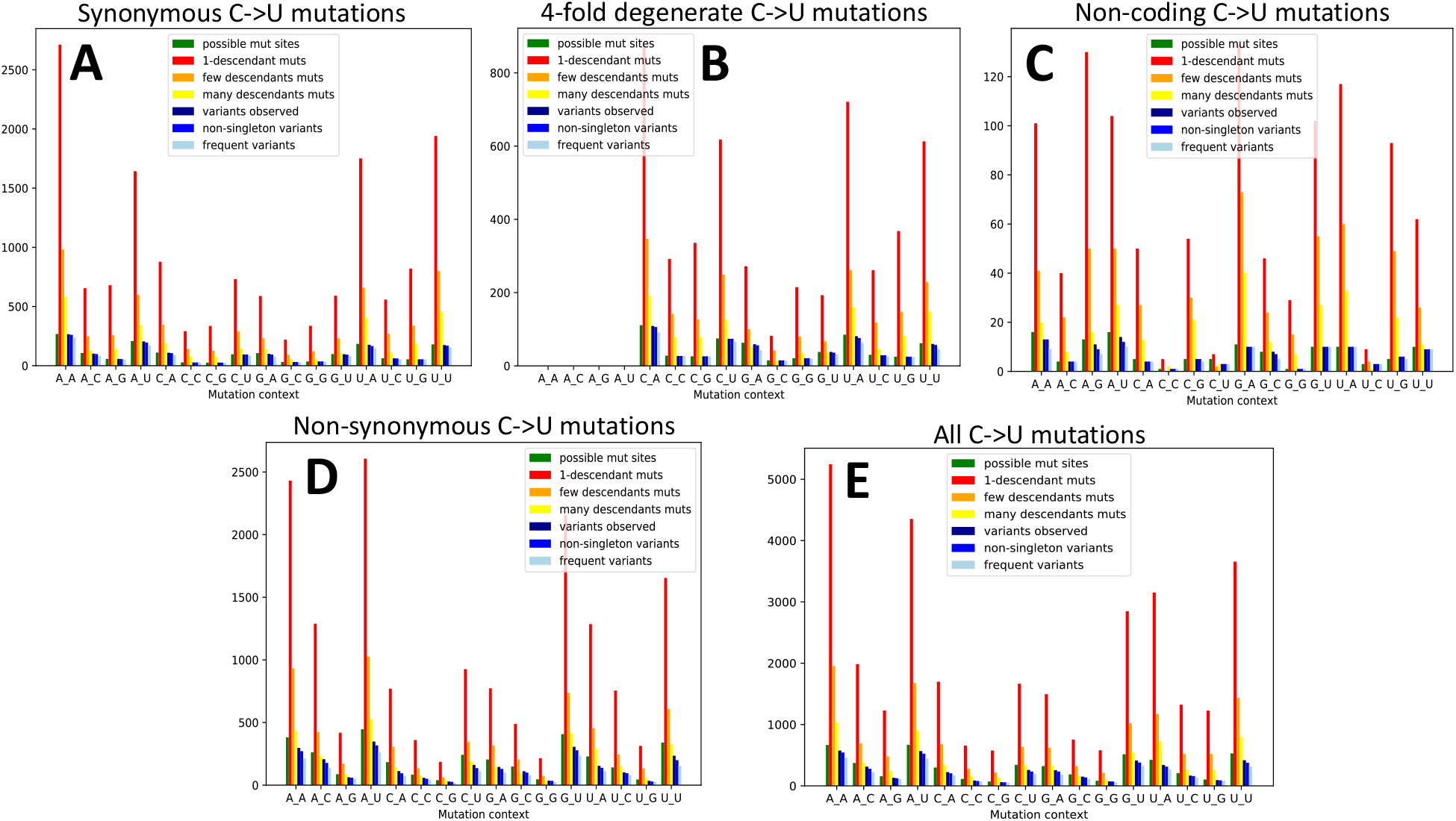
C→U mutation and mutation possibility counts in different base contexts. C→U mutation counts depending on the previous and next base (5’ and 3’ base neighbours, shown on the X axis). A_G represents, for example, the trinucleotide ACG and its mutation counts into trinucleotide AUG. Colors are as in legend Figure 1. **A** synonymous sites, **B** 4-fold degenerate sites, **C** non-coding sites, **D** non-synonymous sites, **E** all sites.

**Figure S6:**
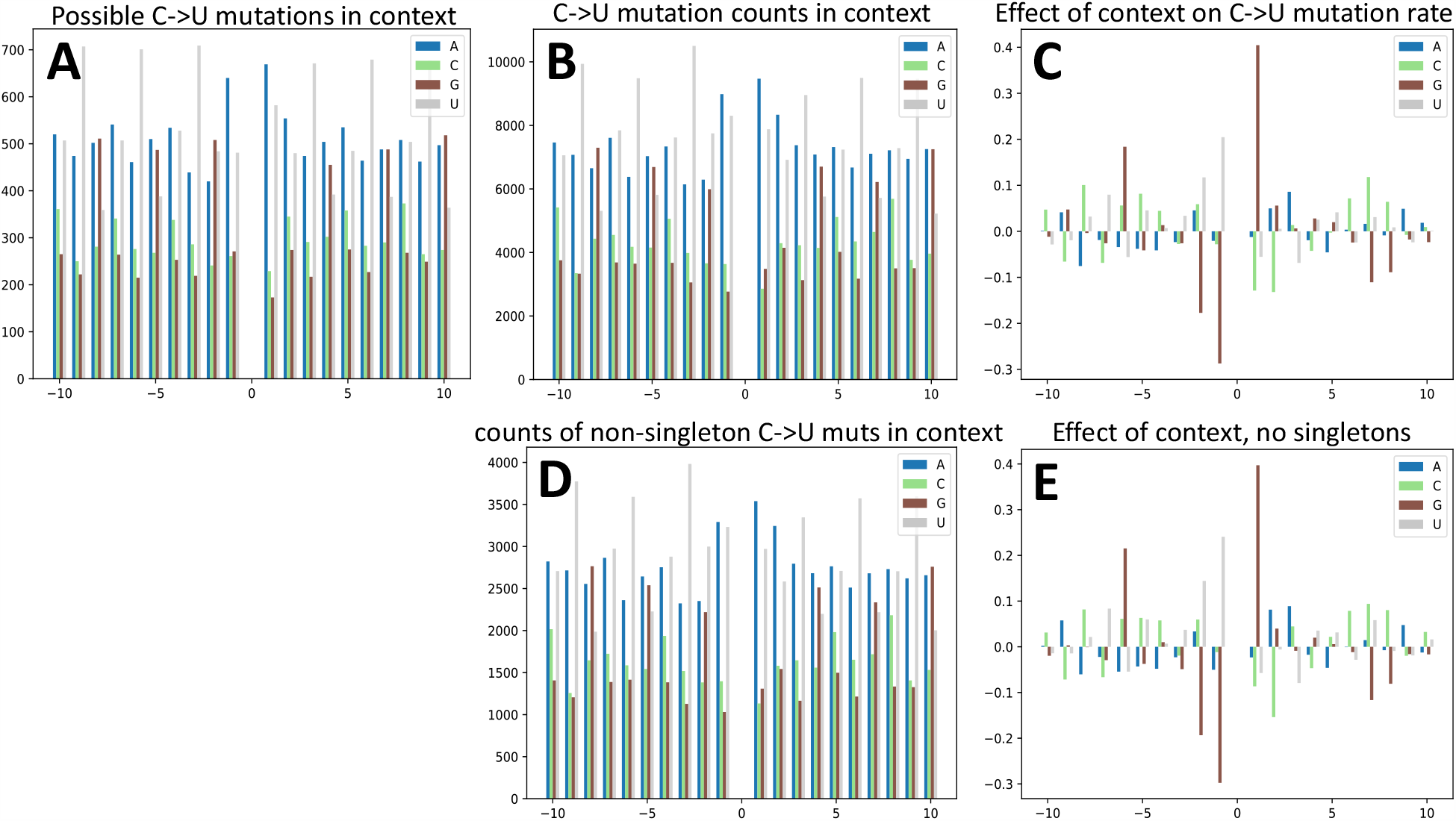
C→U synonymous mutations and mutation rates in different longer-range base contexts. Here we consider only synonymous C→U mutations. X axis values represent the distance of the considered base to the one whose mutation rate is considered. Y axis values represent **A** the numbers of possible synonymous mutations with the given context, **B** the numbers of observed synonymous mutations, **D** the numbers of observed non-singleton mutations, **C** the effect on mutation rate that the considered base at the considered position has, **E** same as **C** but without considering mutations with only one descendant. For example, the value for base G at position -1 in plot **C** represents the increase in GC→GU mutation rate vs all other C→U mutation rates; a Y axis value of 0.1 means that the given context increases the background mutation rate by 10%.

**Figure S7:**
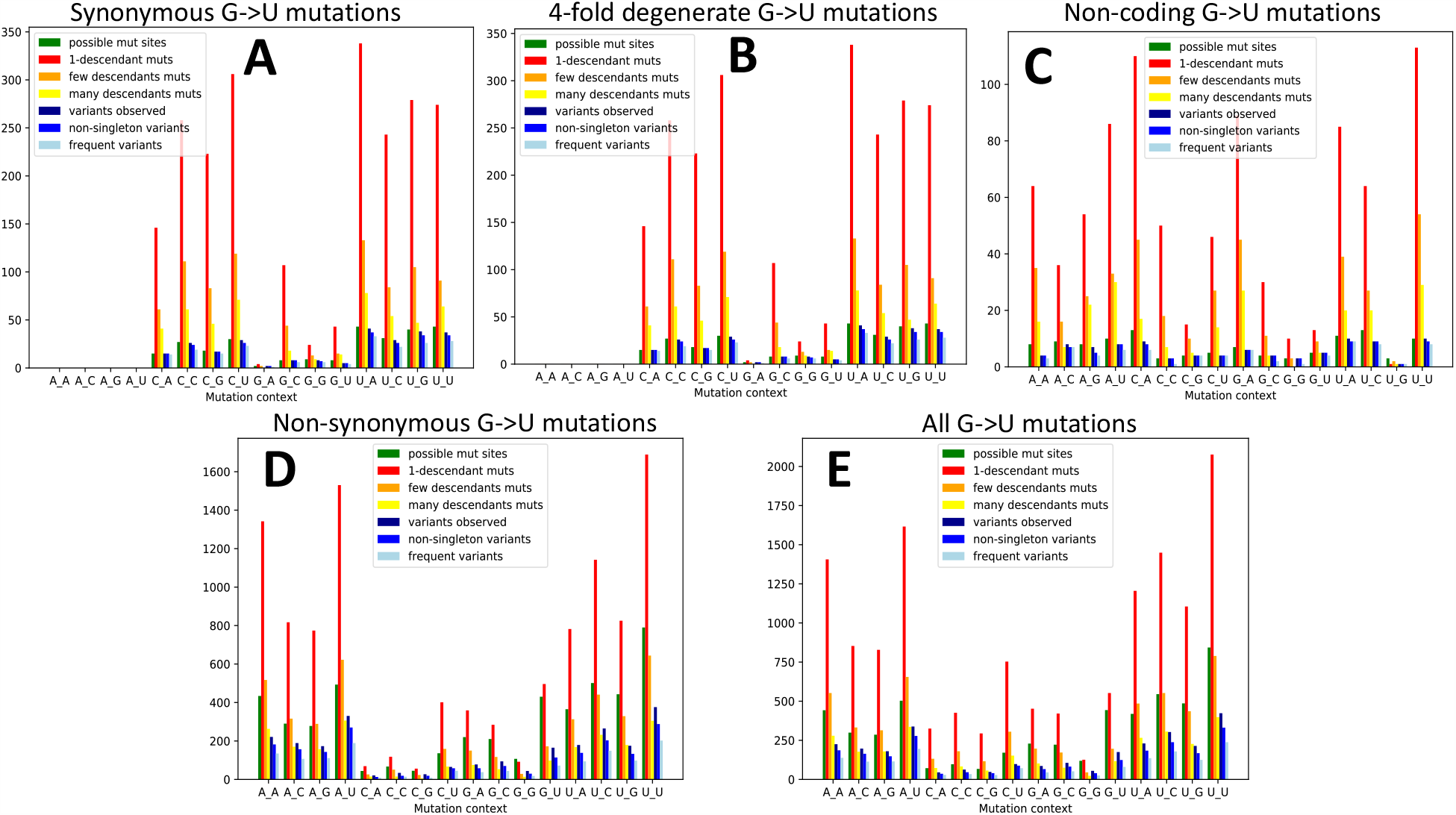
G→U mutation and mutation possibility counts in different base contexts. The X axes show the 16 types of mutation contexts for a G→U mutation, for example C_A means the rate of mutation from trinucleotide CGA to trinucleotide CUA. Colors are as in legends and as in Figure 1. **A** synonymous sites, **B** 4-fold degenerate sites, **C** non-coding sites, **D** non-synonymous sites, **E** all sites.

**Figure S8:**
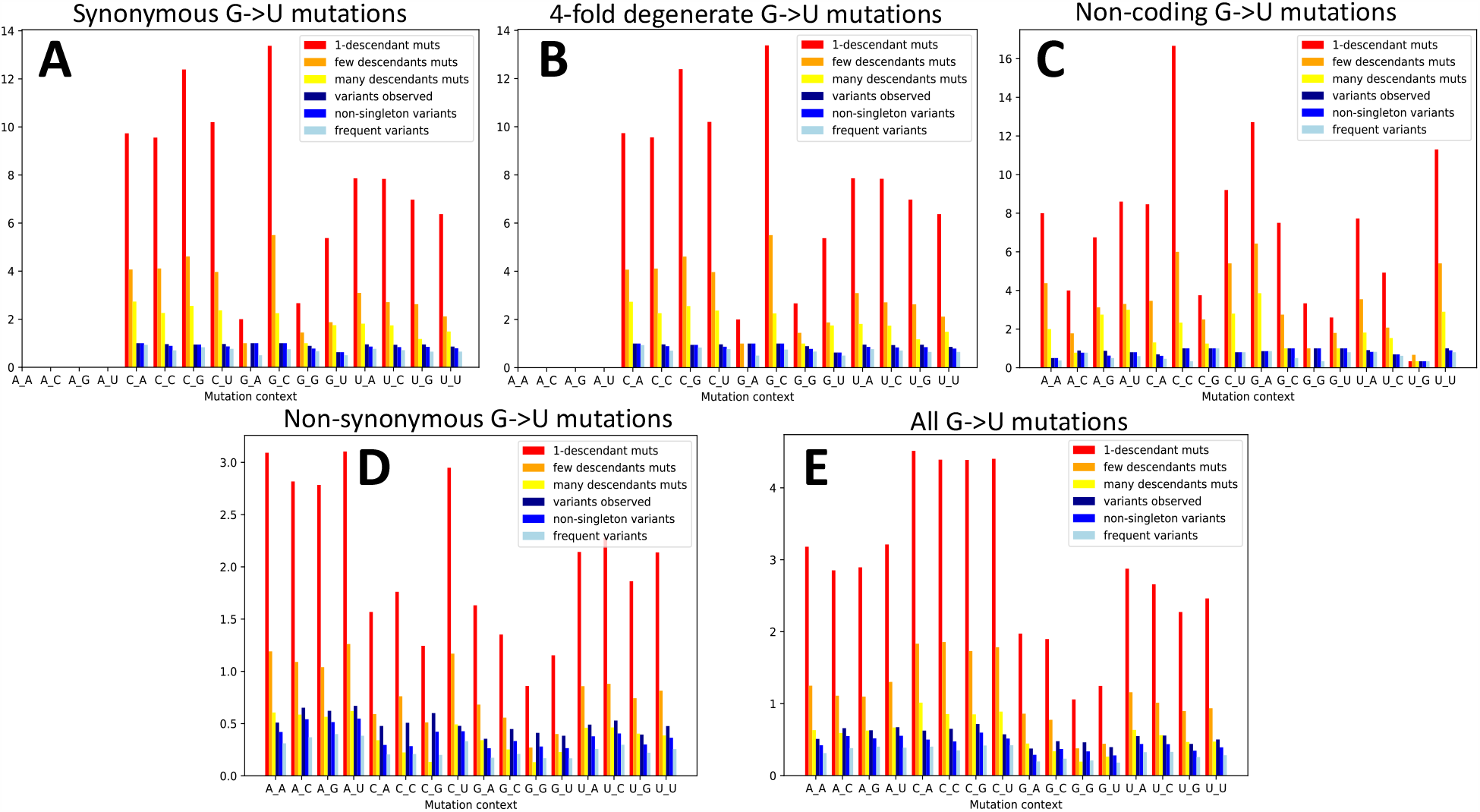
G→U mutation rates in different base contexts. G→U mutation rate depending on the previous and next base (5’ and 3’ base neighbours, shown on the X axis). C_A represents, for example, the trinucleotide CGA and its synonymous mutation rate into trinucleotide CUA. Colors are as in legend Figure 3. **A** synonymous sites, **B** 4-fold degenerate sites, **C** non-coding sites, **D** non-synonymous sites, **E** all sites.

**Figure S9:**
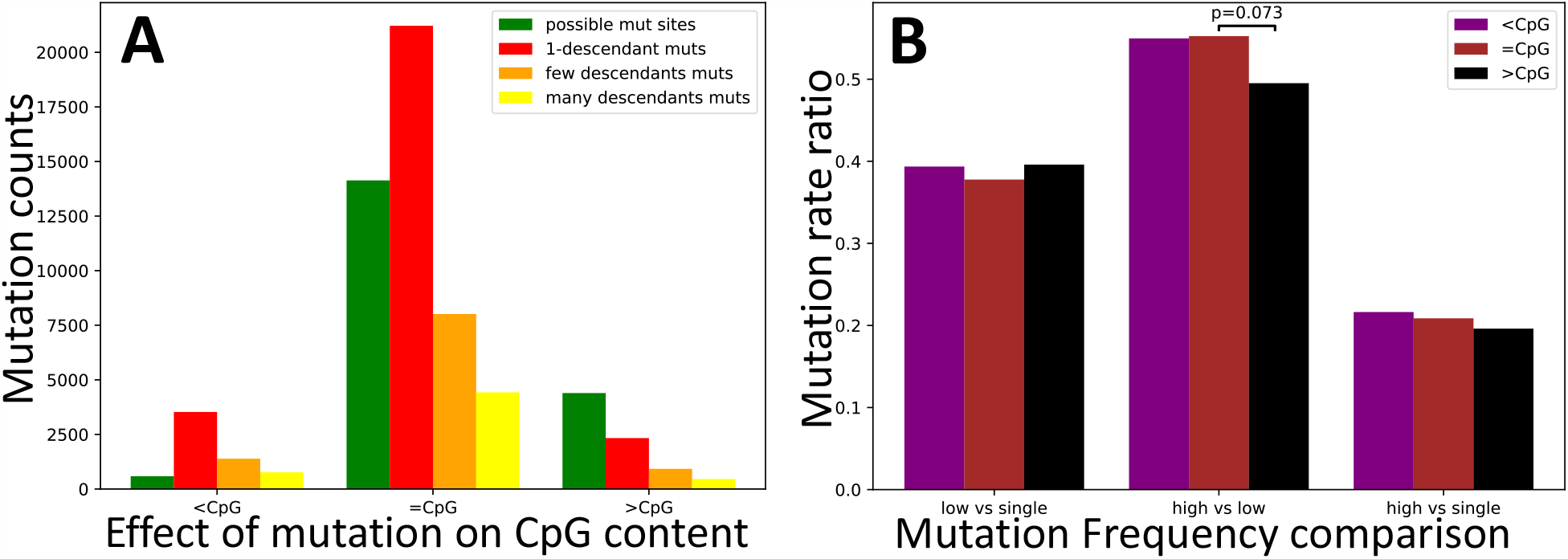
Test of selection affecting CpG content at synonymous sites. Values are the same as in Figure 5, but this time we focus on synonymous mutations that decrease CpG content (“*<*CpG”), increase it (“>CpG”), or leave it unaltered (“=CpG”). Only p-values below 0.1 are shown.

**Figure S10:**
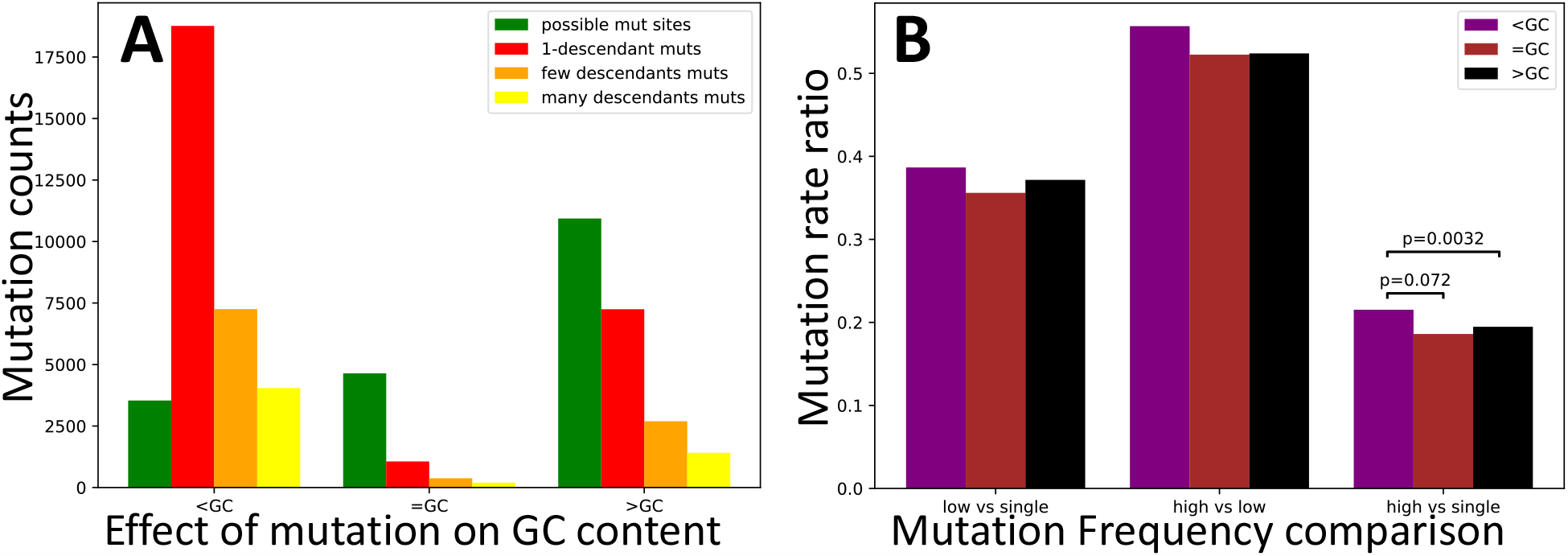
Test of selection affecting GC content at synonymous sites. Values are the same as in Figure 5, but this time we focus on synonymous mutations that decrease GC content (“*<*GC”), increase it (“>GC”), or leave it unaltered (“=GC”). Only p-values below 0.1 are shown.

## References

[1] Yong-Zhen Zhang and Edward C Holmes. A genomic perspective on the origin and emergence of sars-cov-2. Cell, 2020.

[2] Xianding Deng, Wei Gu, Scot Federman, Louis Du Plessis, Oliver Pybus, Nuno Faria, Candace Wang, Guixia Yu, Chao-Yang Pan, Hugo Guevara, et al. A genomic survey of sars-cov-2 reveals multiple introductions into northern california without a predominant lineage. medRxiv, 2020.

[3] Simon Dellicour, Keith Durkin, Samuel L Hong, Bert Vanmechelen, Joan Martí-Carreras, Mandev S Gill, Cécile Meex, Sébastien Bontems, Emmanuel André, Marius Gilbert, et al. A phylodynamic workflow to rapidly gain insights into the dispersal history and dynamics of sars-cov-2 lineages. BioRxiv, 2020.

[4] Fatima Amanat and Florian Krammer. ars-cov-2 vaccines: status report. Immunity, 2020.

[5] Alan M Rice, Atahualpa Castillo Morales, Alexander T Ho, Christine Mordstein, Stefanie Mühlhausen, Samir Watson, Laura Cano, Bethan Young, Grzegorz Kudla, and Laurence D Hurst. Evidence for strong mutation bias towards, and selection against, u content in sars-cov-2: implications for vaccine design. Molecular Biology and Evolution, 2020.

[6] Bethany Dearlove, Eric Lewitus, Hongjun Bai, Yifan Li, Daniel B Reeves, M Gordon Joyce, Paul T Scott, Mihret F Amare, Sandhya Vasan, Nelson L Michael, et al. A sars-cov-2 vaccine candidate would likely match all currently circulating variants. Proceedings of the National Academy of Sciences, 117(38):23652–23662, 2020.

[7] Bette Korber, Will M Fischer, Sandrasegaram Gnanakaran, Hyejin Yoon, James Theiler, Werner Abfalterer, Nick Hengartner, Elena E Giorgi, Tanmoy Bhattacharya, Brian Foley, et al. Tracking changes in sars-cov-2 spike: evidence that d614g increases infectivity of the covid-19 virus. Cell, 182(4):812–827, 2020.

[8] Lucy van Dorp, Damien Richard, Cedric CS Tan, Liam P Shaw, Mislav Acman, and François Balloux. No evidence for increased transmissibility from recurrent mutations in sars-cov-2. bioRxiv, 2020.

[9] Lucy van Dorp, Mislav Acman, Damien Richard, Liam P Shaw, Charlotte E Ford, Louise Ormond, Christopher J Owen, Juanita Pang, Cedric CS Tan, Florencia AT Boshier, et al. Emergence of genomic diversity and recurrent mutations in sars-cov-2. Infection, Genetics and Evolution, page 104351, 2020.

[10] You Li, Ye Wang, Yaping Qiu, Zhen Gong, Lei Deng, Min Pan, Huiping Yang, Jianan Xu, Li Yang, and Jin Li. Sars-cov-2 spike glycoprotein receptor binding domain is subject to negative selection with predicted positive selection mutations. bioRxiv, 2020.

[11] Rachele Cagliani, Diego Forni, Mario Clerici, and Manuela Sironi. Computational inference of selection underlying the evolution of the novel coronavirus, severe acute respiratory syndrome coronavirus 2. Journal of Virology, 94(12), 2020.

[12] Huiguang Yi. 2019 novel coronavirus is undergoing active recombination. Clinical Infectious Diseases, 2020.

[13] Salvatore Di Giorgio, Filippo Martignano, Maria Gabriella Torcia, Giorgio Mattiuz, and Silvestro G Conticello. Evidence for host-dependent rna editing in the transcriptome of sars-cov-2. Science Advances, page eabb5813, 2020.

[14] Alex Graudenzi, Davide Maspero, Fabrizio Angaroni, Rocco Piazza, and Daniele Ramazzotti. Mutational signatures and heterogeneous host response revealed via large-scale characterization of sars-cov-2 genomic diversity. BioRxiv, 2020.

[15] Tobias Mourier, Mukhtar Sadykov, Michael J Carr, Gabriel Gonzalez, William W Hall, and Arnab Pain. Host-directed editing of the sars-cov-2 genome. Biochemical and Biophysical Research Communications, 2020.

[16] Mukhtar Sadykov, Tobias Mourier, Qingtian Guan, and Arnab Pain. Short sequence motif dynamics in the sars-cov-2 genome suggest a role for cytosine deamination in cpg reduction. BioRxiv, 2020.

[17] Sebastian Duchene, Leo Featherstone, Melina Haritopoulou-Sinanidou, Andrew Rambaut, Philippe Lemey, and Guy Baele. Temporal signal and the phylodynamic threshold of sars-cov-2. bioRxiv, 2020.

[18] Jody Phelan, Wouter Deelder, Daniel Ward, Susana Campino, Martin L Hibberd, and Taane G Clark. Controlling the sars-cov-2 outbreak, insights from large scale whole genome sequences generated across the world. BioRxiv, 2020.

[19] Daniele Ramazzotti, Fabrizio Angaroni, Davide Maspero, Carlo Gambacorti-Passerini, Marco Antoniotti, Alex Graudenzi, and Rocco Piazza. Characterization of intra-host sars-cov-2 variants improves phylogenomic reconstruction and may reveal functionally convergent mutations. bioRxiv, 2020.

[20] Rui Wang, Yuta Hozumi, Yong-Hui Zheng, Changchuan Yin, and Guo-Wei Wei. Host immune response driving sars-cov-2 evolution. Viruses, 12(10):1095, 2020.

[21] Peter Simmonds. Rampant c-> u hypermutation in the genomes of sars-cov-2 and other coronaviruses–causes and consequences for their short and long evolutionary trajectories. bioRxiv, 2020.

[22] Nicola De Maio, Conor Walker, Rui Borges, Lukas Weilguny, Greg Slodkowicz, and Nick Goldman. Issues with sars-cov-2 sequencing data, 2020.

[23] Yatish Turakhia, Nicola De Maio, Bryan Thornlow, Landen Gozashti, Robert Lanfear, Conor R. Walker, Angie S. Hinrichs, Jason D. Fernandes, Rui Borges, Greg Slodkowicz, Lukas Weilguny, David Haussler, Nick Goldman, and Russell Corbett-Detig. Stability of SARS-CoV-2 phylogenies. PLOS Genetics, 16(11):e1009175, November 2020.

[24] Ziheng Yang. Paml 4: phylogenetic analysis by maximum likelihood. Molecular biology and evolution, 24(8):1586–1591, 2007.

[25] Sergei L Kosakovsky Pond and Spencer V Muse. Hyphy: hypothesis testing using phylogenies. In Statistical methods in molecular evolution, pages 125–181. Springer, 2005.

[26] José M Cuevas, Pilar Domingo-Calap, and Rafael Sanjuán. The fitness effects of synonymous mutations in dna and rna viruses. Molecular biology and evolution, 29(1):17–20, 2012.

[27] John H McDonald and Martin Kreitman. Adaptive protein evolution at the adh locus in drosophila. Nature, 351(6328):652–654, 1991.

[28] Stephen J Freeland and Laurence D Hurst. The genetic code is one in a million. Journal of molecular evolution, 47(3):238–248, 1998.

[29] Nick Goldman. Further results on error minimization in the genetic code. Journal of molecular evolution, 37(6):662–664, 1993.

[30] Hagit T Porath, Shai Carmi, and Erez Y Levanon. A genome-wide map of hyper-edited rna reveals numerous new sites. Nature communications, 5(1):1–10, 2014.

[31] Nga Ling Ko, Emmanuel Birlouez, Simon Wain-Hobson, Renaud Mahieux, and Jean-Pierre Vartanian. Hyper-editing of human t-cell leukemia virus type 2 and simian t-cell leukemia virus type 3 by the dsrna adenosine deaminase adar-1. The Journal of General Virology, 93(Pt 12):2646, 2012.

[32] Masato Kosuge, Emi Furusawa-Nishii, Koyu Ito, Yoshiro Saito, and Kouetsu Ogasawara. Point mutation bias in sars-cov-2 variants results in increased ability to stimulate inflammatory responses. Scientific reports, 10(1):1–9, 2020.

[33] Patrick CY Woo, Beatrice HL Wong, Yi Huang, Susanna KP Lau, and Kwok-Yung Yuen. Cytosine deamination and selection of cpg suppressed clones are the two major independent biological forces that shape codon usage bias in coronaviruses. Virology, 369(2):431–442, 2007.

[34] Alexander Y Panchin and Yuri V Panchin. Excessive g–u transversions in novel allele variants in sars-cov-2 genomes. PeerJ, 8:e9648, 2020.

[35] Nicolae Sapoval, Medhat Mahmoud, Michael D Jochum, Yunxi Liu, RA Leo Elworth, Qi Wang, Dreycey Albin, Huw Ogilvie, Michael D Lee, Sonia Villapol, et al. Hidden genomic diversity of sars-cov-2: implications for qrt-pcr diagnostics and transmission. BioRxiv, 2020.

[36] Pauli Virtanen, Ralf Gommers, Travis E Oliphant, Matt Haberland, Tyler Reddy, David Cournapeau, Evgeni Burovski, Pearu Peterson, Warren Weckesser, Jonathan Bright, et al. Scipy 1.0: fundamental algorithms for scientific computing in python. Nature methods, 17(3):261–272, 2020.

[37] Thomas H Jukes, Charles R Cantor, et al. Evolution of protein molecules. Mammalian protein metabolism, 3:21–132, 1969.

[38] Masami Hasegawa, Hirohisa Kishino, and Taka-aki Yano. Dating of the human-ape splitting by a molecular clock of mitochondrial dna. Journal of molecular evolution, 22(2):160–174, 1985.

[39] Nick Goldman and Ziheng Yang. A codon-based model of nucleotide substitution for protein-coding dna sequences. Molecular biology and evolution, 11(5):725–736, 1994.

[40] Ziheng Yang. Estimating the pattern of nucleotide substitution. Journal of molecular evolution, 39(1):105–111, 1994.

[41] Bui Quang Minh, Heiko A Schmidt, Olga Chernomor, Dominik Schrempf, Michael D Woodhams, Arndt Von Haeseler, and Robert Lanfear. Iq-tree 2: New models and efficient methods for phylogenetic inference in the genomic era. Molecular Biology and Evolution, 37(5):1530–1534, 2020.

[42] Alexey M Kozlov, Diego Darriba, Tomáš Flouri, Benoit Morel, and Alexandros Stamatakis. Raxml-ng: a fast, scalable and user-friendly tool for maximum likelihood phylogenetic inference. Bioinformatics, 35(21):4453–4455, 2019.

[43] Benoit Morel, Pierre Barbera, Lucas Czech, Ben Bettisworth Lukas Hübner, Sarah Lutteropp, Dora Serdari, Evangelia-Georgia Kostaki, Ioannis Mamais, Alexey Kozlov, et al. Phylogenetic analysis of sars-cov-2 data is difficult. bioRxiv, 2020.

[44] Morgan N Price, Paramvir S Dehal, and Adam P Arkin. Fasttree 2–approximately maximum-likelihood trees for large alignments. PloS one, 5(3):e9490, 2010.

[45] Jack Kuipers, Aashil A Batavia, Kim Philipp Jablonski, Fritz Bayer, Nico Borgsmüller, Arthur Dondi, Monica-Andreea Drăgan, Pedro Ferreira, Katharina Jahn, Lisa Lamberti, et al. Within-patient genetic diversity of sars-cov-2. bioRxiv, 2020.

[46] Daniel J Wilson, Ryan D Hernandez, Peter Andolfatto, and Molly Przeworski. A population genetics-phylogenetics approach to inferring natural selection in coding sequences. PLoS Genet, 7(12):e1002395, 2011.

[47] Nicola De Maio, Christian Schlötterer, and Carolin Kosiol. Linking great apes genome evolution across time scales using polymorphism-aware phylogenetic models. Molecular biology and evolution, 30(10):2249–2262, 2013.

[48] Kai Zeng and Brian Charlesworth. Estimating selection intensity on synonymous codon usage in a nonequilibrium population. Genetics, 183(2):651–662, 2009.

[49] F Clemente and C Vogl. Evidence for complex selection on four-fold degenerate sites in drosophila melanogaster. Journal of evolutionary biology, 25(12):2582–2595, 2012.

[50] Steven Kemp, William Harvey, Rawlings Datir, Dami Collier, Isabella Ferreira, Alessandro Carabelii, David L Robertson, and Ravindra K Gupta. Recurrent emergence and transmission of a sars-cov-2 spike deletion δh69/v70. bioRxiv, 2020.

[51] Rob Lanfear. A global phylogeny of sars-cov-2 sequences from gisaid., 2020.

[52] Kazutaka Katoh and Daron M Standley. Mafft multiple sequence alignment software version 7: improvements in performance and usability. Molecular biology and evolution, 30(4):772–780, 2013.

[53] Ole Tange et al. Gnu parallel-the command-line power tool. The USENIX Magazine, 36(1):42–47, 2011.

[54] Thomas Junier and Evgeny M Zdobnov. The newick utilities: high-throughput phylogenetic tree processing in the unix shell. Bioinformatics, 26(13):1669–1670, 2010.

[55] Uyen Mai and Siavash Mirarab. Treeshrink: fast and accurate detection of outlier long branches in collections of phylogenetic trees. BMC genomics, 19(5):23–40, 2018.

[56] Yatish Turakhia, Bryan Thornlow, Angie S Hinrichs, Nicola De Maio, Landen Gozashti, Robert Lanfear, David Haussler, and Russ Corbett-Detig. Ultrafast sample placement on existing trees (usher) empowers real-time phylogenetics for the sars-cov-2 pandemic. bioRxiv, 2020.

